# Hybridisation capture allows DNA damage analysis of ancient marine eukaryotes

**DOI:** 10.1101/2020.09.25.310920

**Authors:** L. Armbrecht, G. Hallegraeff, C.J.S. Bolch, C. Woodward, A. Cooper

## Abstract

Marine sedimentary ancient DNA (*sed*aDNA) is increasingly used to study past ocean ecosystems, however, studies have been severely limited by the very low amounts of DNA preserved in the subseafloor, and the lack of bioinformatic tools to authenticate *sed*aDNA in metagenomic data. We applied a hybridisation capture ‘baits’ technique to target marine eukaryote *sed*aDNA (specifically, phytoplankton, ‘Phytobaits1’; and harmful algal bloom taxa, ‘HABbaits1’), which resulted in up to 4- and 9-fold increases, respectively, in the relative abundance of eukaryotes compared to shotgun sequencing. We further used the new bioinformatic tool ‘HOPS’ to authenticate the *sed*aDNA component, establishing a new proxy to assess *sed*aDNA authenticity, the Ancient: Default (A:D) sequences ratio, here positively correlated with subseafloor depth, and generated the first-ever DNA damage profiles of a key phytoplankton, the ubiquitous coccolithophore *Emiliania huxleyi.* Our study opens new options for the detailed investigation of marine eukaryotes and their evolution over geological timescales.

## 1 Introduction

Over the past decade marine sedimentary ancient DNA (*sed*aDNA) has become increasingly used to study past ocean ecosystems and oceanographic conditions. The novelty of using *sed*aDNA lies in its enormous potential to detect genetic signals of taxa that do and don’t fossilise – meaning that in theory it is possible to go beyond standard environmental proxies and facilitate reconstruction of past marine ecosystems across the entire food web. For example, *sed*aDNA has revealed relationships between past marine community composition and paleo-tsunami episodes in Japan over the past 2,000 years (Szczuciński et al., 2016), oxygen minimum zone expansions in the temperate Arabian Sea region over 43 thousand years (kyr) (More et al., 2018), and Arctic sea-ice conditions spanning 100kyr (DeSchepper et al., 2019). While the logistical challenge of acquiring undisturbed sediment cores from the deep seafloor remains, the field of *sed*aDNA research is rapidly advancing due to new ship-board core sampling procedures that allow far greater contamination control, and improvements in sample processing, sequencing technologies and bioinformatic tools (Armbrecht et al., 2019).

Among the huge diversity of marine eukaryotes, phytoplankton are particularly useful targets to study past ocean conditions. Phytoplankton are free-floating, unicellular microalgae fulfilling two important functions: *(1)* they form the base of the marine food web supporting virtually all higher trophic organisms (*e.g.,* Verity and Smetacek, 1990), and (*2)* are highly useful environmental indicators due to their sensitivity to changing physical and chemical oceanographic conditions (Hays et al., 2005). After phytoplankton die, they sink to the seafloor where small proportions of their DNA are able to become entombed and preserved in sediments under favorable conditions, over time forming long-term records of past ocean and climate conditions. Using the small subunit ribosomal RNA gene (18S rRNA, a common taxonomic marker gene), we recently determined the fraction of marine eukaryote *sed*aDNA preserved in Tasmanian coastal sediments to be a mere 1.37% of the total *sed*aDNA pool (Armbrecht et al., 2020). A slightly higher proportion of eukaryote *sed*aDNA (and also higher diversity) may be captured by combining multiple taxonomic markers, e.g., the small and large subunit ribosomal RNA gene (Armbrecht, 2020). However, rather than analysing only part of the total *sed*aDNA pool (such as eukaryote marker genes within a large metagenomic dataset), it would be much more cost-effective to increase marine eukaryote *sed*aDNA yield by optimising extraction and laboratory protocols, to maximise sequencing of *sed*aDNA from the intended target organisms.

Metagenomic approaches extract and analyse the ‘total’ DNA in a sample (‘shotgun’ style), irrespective of the source organism, facilitating recovery of DNA sequences from any organism in proportion to their original presence in that sample. As a result, metagenomic approaches are well suited to the study of microbial and environmental ancient DNA (e.g., Taberlet et al., 2012; Pedersen et al., 2015; Weyrich et al., 2017), including *sed*aDNA. The use of metagenomics does not prescribe the target DNA fragment size and preserve DNA damage patterns characteristic of ancient DNA. Importantly, the combination of DNA fragment size variability and damage patterns are vital to assess the authenticity of potential ancient genetic signals.

Hybridisation capture techniques are an increasingly popular method to focus the metagenomic analysis towards loci of interest, such as specific sequences to investigate particular groups of organisms (Horn et al., 2012; Foster et al., 2020). Hybridisation capture uses short RNA probes (also called ‘baits’) designed to be complementary to DNA sequences of interest (*e.g.*, taxonomic marker genes; Fig. 1). By binding to the target sequence, these genetic baits ‘capture’ DNA fragments from DNA extracts in a manner that preserves size variability, along with DNA-damage patterns that can be used to examine whether sequences appear ancient. Additionally, careful bait design (i.e., selection of target sequences) and optimisations of the application protocol (e.g., hybridisation-temperature settings) allow differing levels of specificity in the capture process. While such ‘baits’ approaches have previously been used to investigate human, animal and even environmental DNA (Paijmans et al., 2013; Li et al., 2015; Murchie et al., 2020), its application to marine sediments to capture *sed*aDNA from key primary producers and environmental indicator organisms (e.g., eukaryotic phytoplankton) remains untested.

**Figure 1:**
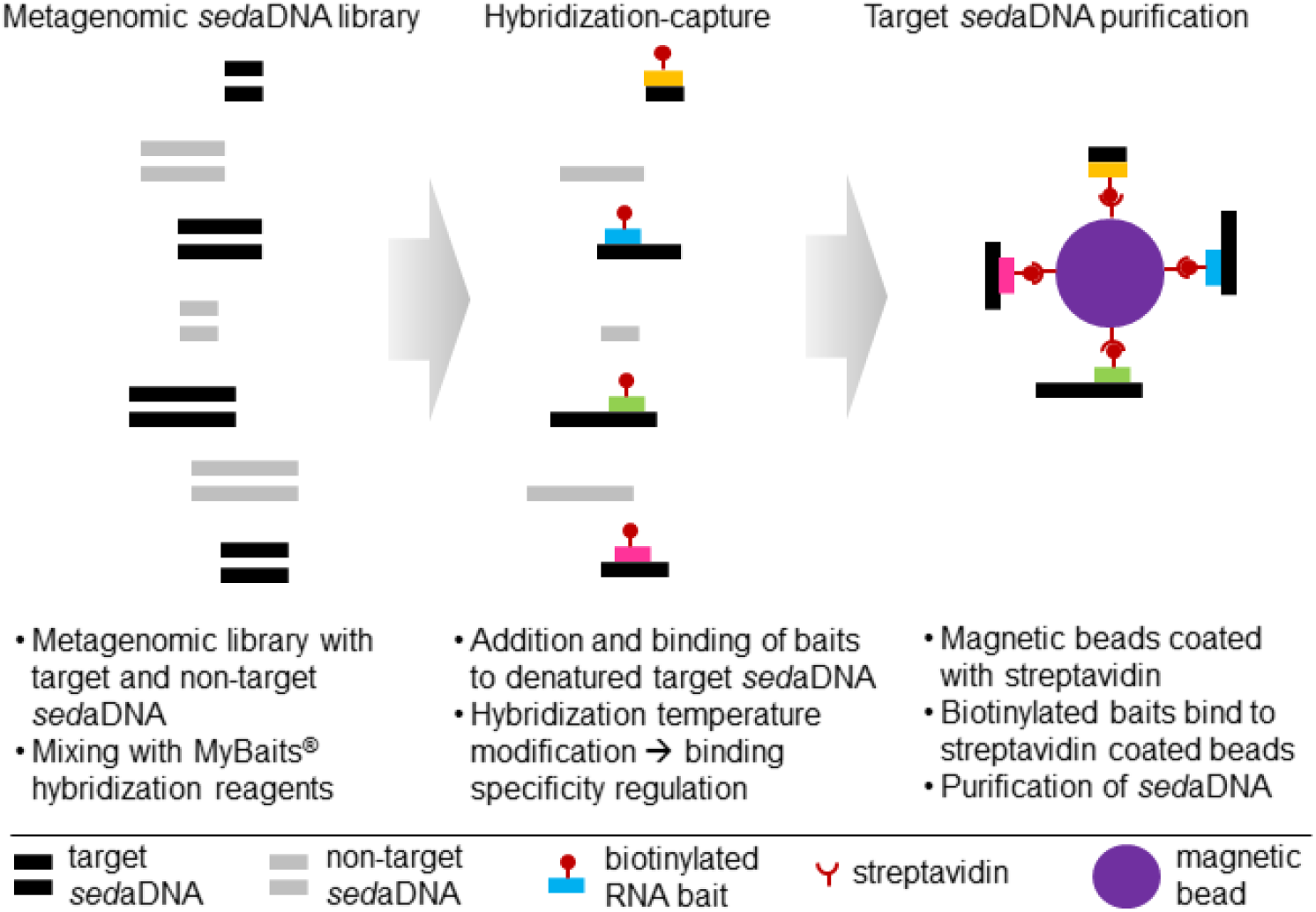
Schematic of hybridisation capture applied to marine sedimentary ancient DNA (*sed*aDNA). The three main steps are the preparation of a metagenomic *sed*aDNA library, hybridisation capture using RNA baits (in this study: Phytobaits1 and HABbaits1) that are biotinylated, which enables binding of baits to streptavidin-coated magnetic beads (multiple baits per bead possible, schematic not to scale). For further technical details see Methods, MyBaits® Manual V4.01 (2018), and Horn et al. (2012).

The assessment of *sed*aDNA authenticity has been hindered by a lack of established approaches to identify and analyse DNA damage patterns of rare ancient microorganisms in metagenomic samples (such as eukaryotes in marine *sed*aDNA). For example, software commonly used to detect DNA damage patterns, such as ‘mapDamage’, computes nucleotide misincorporation and fragmentation patterns by mapping next-generation sequencing reads against a reference genome (Ginolhac et al., 2012; Jónsson et al., 2013). This requires high-quality modern reference genomes, or species where ancient DNA is available in sufficient quantity (e.g., animals or humans; Llamas et al., 2015; Tobler et al., 2017), but neither is generally possible with marine eukaryote *sed*aDNA. There is a lack of high-quality reference sequences for the thousands of marine organisms occurring in the global ocean, and the threshold of ~250 reads per species required to analyse and plot DNA damage patterns in mapDamage (Collin et al., 2020) is often not reached in *sed*aDNA. Recently, Hübler et al. (2020) developed a new bioinformatic tool HOPS - ‘Heuristic Operations for Pathogen Screening’ - based on the mapDamage algorithm, to identify and authenticate bacterial pathogens in ancient metagenomic samples and extract this information for further downstream analysis. In combination with hybridisation capture to generate a larger number of ancient eukaryote sequences, HOPS has the potential to allow the assessment of *sed*aDNA authenticity based on DNA damage profiles from key marine eukaryotes, even if only very few sequences are available (>50 reads per species, Hübler et al., 2020).

Here, we develop and apply two hybridisation capture bait sets for the first such analysis of marine sediments, targeting (*i*) marine phytoplankton very broadly for general paleo-monitoring (Phytobaits1), and (*ii*) selected microalgae (including key phytoplankton groups such as diatoms, dinoflagellates and coccolithophores) that are highly abundant and/or the cause of harmful algal blooms (HABs) in our study region of the East Australian coast (HABbaits1). Based on samples from two coastal sediment cores collected near Maria Island, Tasmania, we demonstrate: 1) the suitability of Phytobaits1 and HABbaits1 as effective tools to maximise *sed*aDNA originating from eukaryote targets relative to shotgun data; 2) the authenticity of both shotgun- and baits-derived sequencing data via HOPS; 3) examine relationships between the ‘ancient’ DNA fraction and subseafloor depth through the development of a new *sed*aDNA proxy; and 4) generate the first-ever DNA damage profile for a keystone marine phytoplankton, the coccolithophore *Emiliania huxleyi.*

## 2 Methods

### 2.1 Samples

Cores were collected during the *RV Investigator* voyage IN2018_T02 (19 and 20 May 2018, respectively, Fig. 2) to Tasmania, from sites in the Mercury Passage and Maria Island (Fig. 2). We collected one KC Denmark Multi-Core (MCS3, 36 cm long, estimated to cover the last ~135 years based on ^210^Pb dating at the Australian Nuclear Science and Technology Organisation (ANSTO, Lucas Heights, Sydney) in the Mercury Passage (MP, 42.550 S, 148.014 E; 68 m water depth), and one gravity core (GC2; 3 m long) offshore from Maria Island (42.845 S, 148.240 E; 104 m) composed of 2 sections; GC2A (bottom) and GC2B (top) estimated to cover the last ~8,000 years based on ^14^C dating, ANSTO). The untreated cores were initially stored on-board at 10°C, followed by transport to and storage at 4 °C at ANSTO. To minimise contamination during core slicing and subsampling (October, 2018, ANSTO), we wiped working benches, sampling and cutting tools with bleach and 80% EtOH, changed gloves immediately when contaminated with sediment, and wore appropriate PPE at all times (gloves, facemask, hairnet, disposable lab gown). We removed the outer ~1 cm of the working core-half (working from bottom to the top of the core), then collected plunge samples by pressing sterile 15 mL centrifuge tubes (Falcon) ~2 cm deep into the sediment core centre at 5 cm depth intervals. All *sed*aDNA samples were immediately frozen at −20°C and transported to the Australian Centre for Ancient DNA (ACAD), Adelaide. For this study, a total of 30 samples were selected from both cores, representing ~2 cm depth intervals within the upper 36 cm of MCS3 and GC2, and ~20 cm depth intervals in GC2 downcore from 36 cm below seafloor (cmbsf).

**Figure 2:**
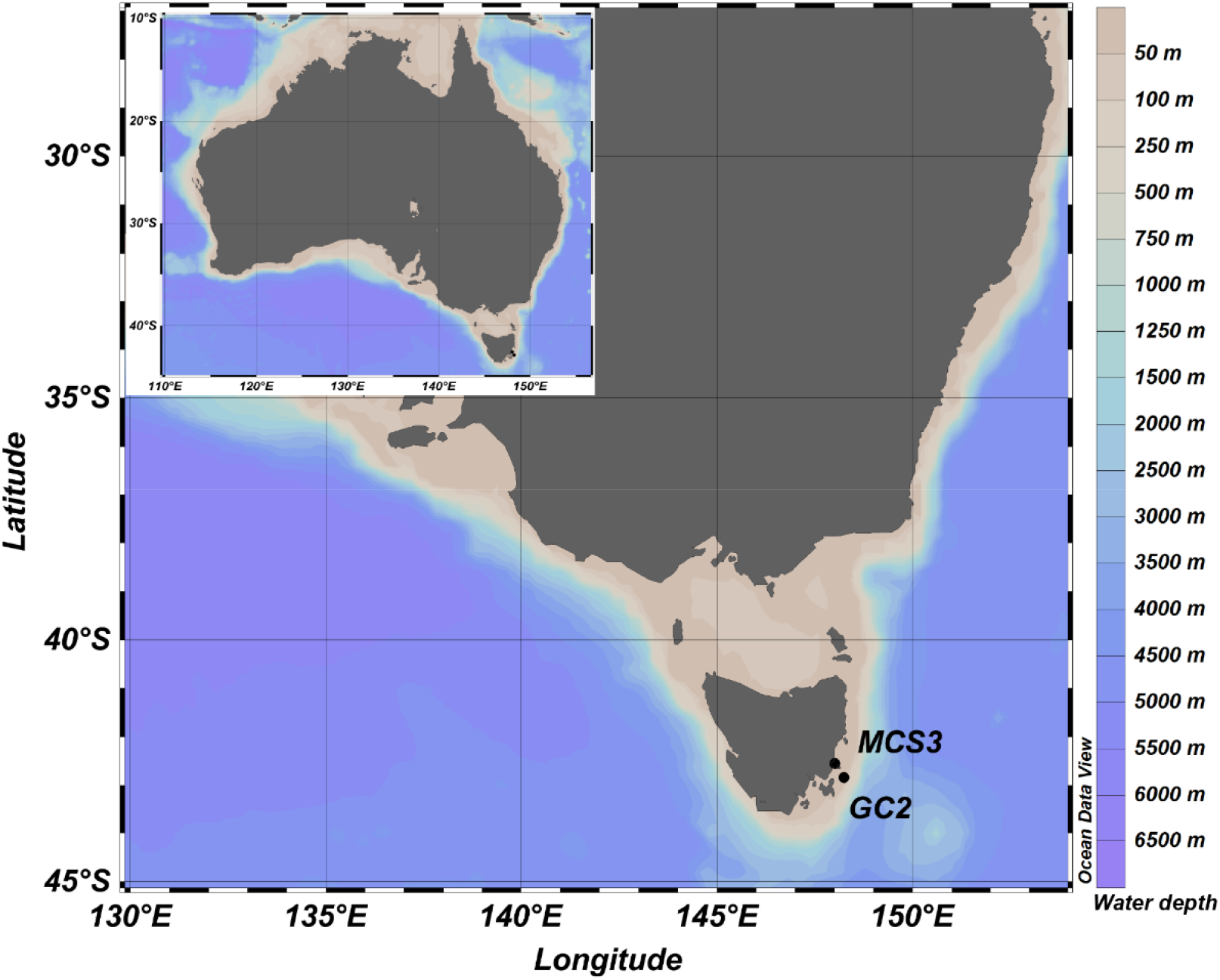
Map of coring sites, inshore (MCS3) and offshore (GC2B) of Maria Island, Tasmania, South-East Australian Coast. Map created in ODV (Schlitzer, R., Ocean Data View, https://odv.awi.de, 2018).

### 2.2 SedaDNA extractions

We prepared *sed*aDNA extracts and sequencing libraries at ACAD’s ultra-clean ancient (GC2) and forensic (MCS3) facilities following ancient DNA decontamination standards (Willerslev and Cooper, 2005). All sample tubes were wiped with bleach on the outside prior to entering the laboratory for subsampling. Our extraction method followed the optimised (“combined”) approach outlined in detail in Armbrecht et al. (2020), with a minor modification in that we stored the final purified DNA in TLE buffer (50 μL Tris HCL (1M), 10 μL EDTA (0.5M), 5 mL nuclease-free water) instead of customary Elution Buffer (Qiagen) (see Supplementary Material Methods). To monitor laboratory contamination, we used extraction blank controls (EBCs) by processing 1-2 (depending on the extraction-batch size) empty bead-tubes through the extraction protocol. A total of 30 extracts were generated from sediment samples and 7 extracts from EBCs.

### 2.3 RNA-baits design

We designed two RNA hybridisation bait-sets, one targeting phytoplankton for a more detailed overview of phytoplankton diversity (hereafter ‘Phytobaits1’), and one targeting specific plankton organisms and their predators to enable detailed investigation of HABs, especially those caused by dinoflagellates, in coastal marine ecosystems (hereafter, ‘HABbaits1’). Phytobaits1 was based on 18S-V9 and 16S-V4 sequences of major phyto- and zooplankton groups, whereas we designed HABbaits1 from a collection of LSU, SSU, D1-D2-LSU, COI, rbcL and ITS sequences for specific marine target organisms often associated with HABs in our study region (Table 1; Supplementary Material Methods). In collaboration with Arbor Biosciences, USA, we designed RNA baits based on these target sequences, with Phytobaits1 containing a total of 15,952 RNA baits targeting the 18S-V9 region of a broad diversity of phytoplankton and their predators and the 16S-V4 region of three cyanobacteria, and HABbaits1 contained 15,310 RNA baits targeting commercially important toxic microalgae and their predators (see Supplementary Material Methods).

**Table 1:**
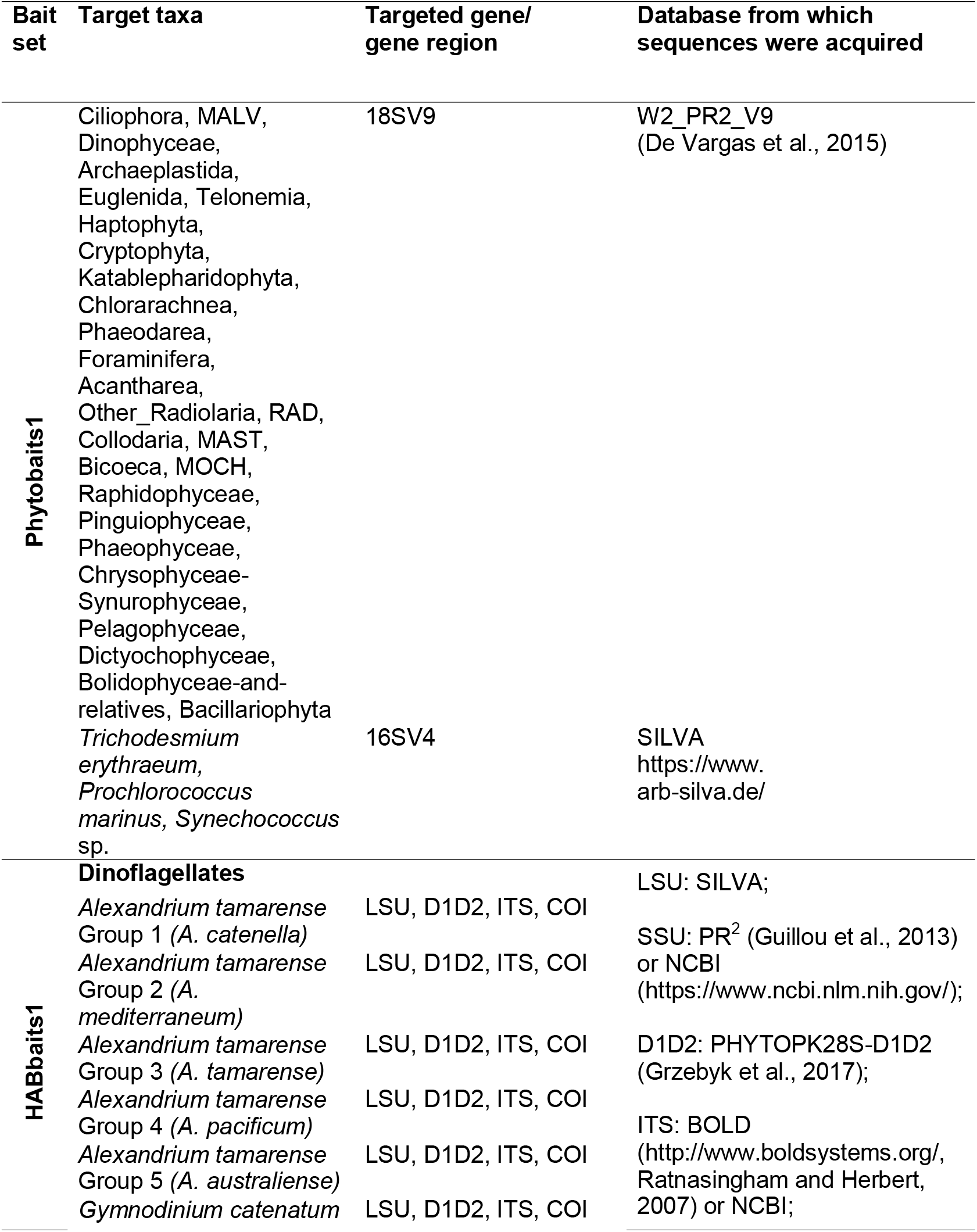
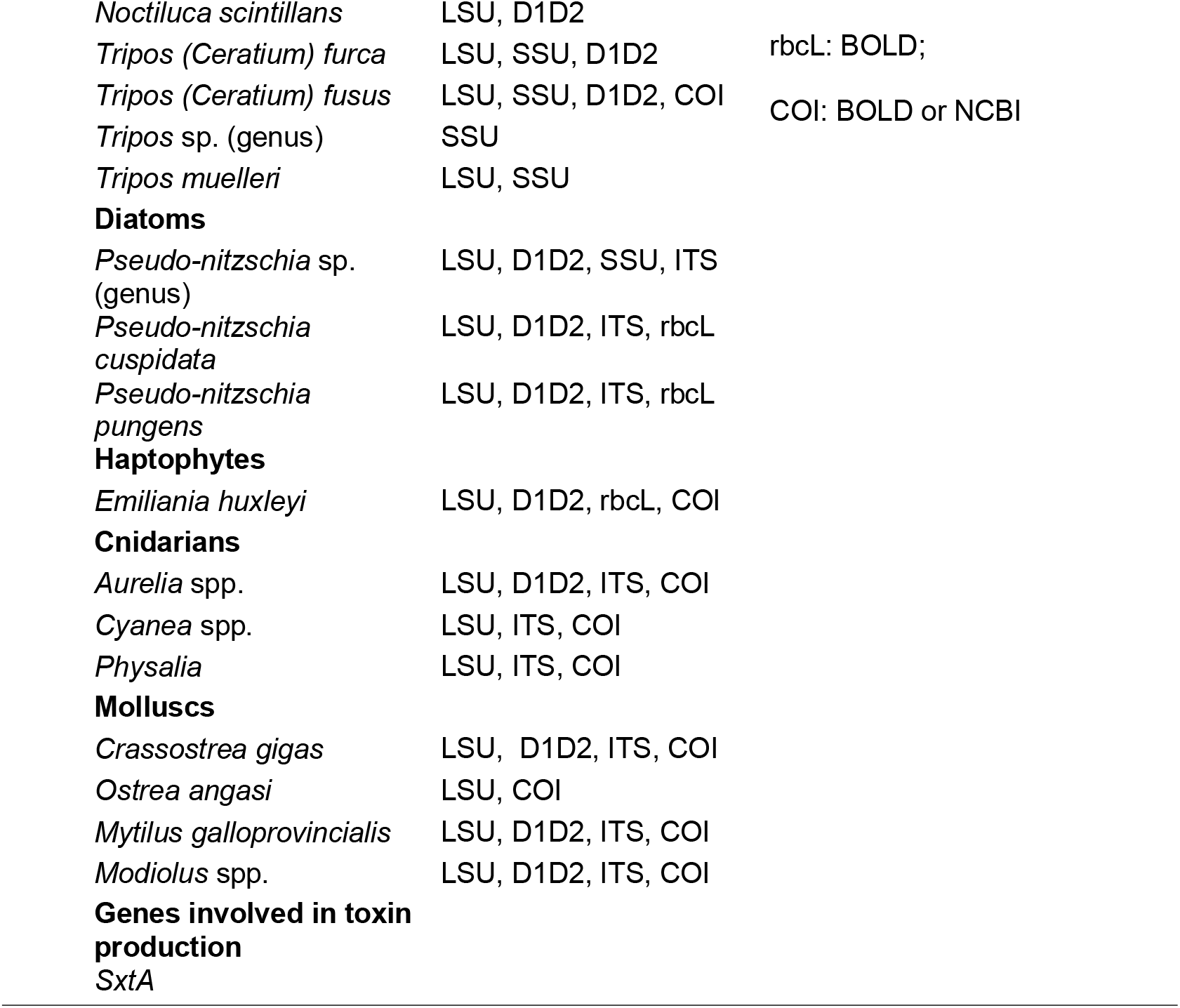
Phytobaits1 and HABbaits1. Target taxa of Phyto- and HABbaits1 genes/gene regions and source databases. For HABbaits1, all listed databases were searched for each gene (region) per target taxon, and, if available, the longest sequence was selected and included.

| Bait set | Target taxa | Targeted gene/ gene region | Database from which sequences were acquired |
| --- | --- | --- | --- |
| Phytobaits1 | Ciliophora, MALV, Dinophyceae, Archaeplastida, Euglenida, Telonemia, Haptophyta, Cryptophyta, Katablepharidophyta, Chlorarachnea, Phaeodarea, Foraminifera, Acantharea, Other_Radiolaria, RAD, Collodaria, MAST, Bicoeca, MOCH, Raphidophyceae, Pinguiphyceae, Phaeophyceae, Chrysophyceae-Synurophyceae, Pelagophyceae, Dictyochophyceae, Bolidophyceae-and-relatives, Bacillariophyta | 18SV9 | W2_PR2_V9 (De Vargas et al., 2015) |
|  | *Trichodesmium erythraeum*, *Prochlorococcus marinus*, *Synechococcus* sp. | 16SV4 | SILVA<br>https://www.arb-silva.de/ |
| HABbaits1 | **Dinoflagellates** |  | LSU: SILVA; |
|  | *Alexandrium tamarens* Group 1 (*A. catenella*) | LSU, D1D2, ITS, COI | SSU: PR<sup>2</sup> (Guillou et al., 2013) or NCBI (https://www.ncbi.nlm.nih.gov/); |
|  | *Alexandrium tamarens* Group 2 (*A. mediterraneum*) | LSU, D1D2, ITS, COI | D1D2: PHYTOPK28S-D1D2 (Grzebyk et al., 2017); |
|  | *Alexandrium tamarens* Group 3 (*A. tamarens*) | LSU, D1D2, ITS, COI | ITS: BOLD (http://www.boldsystems.org/, Ratnasingham and Herbert, 2007) or NCBI; |
|  | *Alexandrium tamarens* Group 4 (*A. pacificum*) | LSU, D1D2, ITS, COI |  |
|  | *Alexandrium tamarens* Group 5 (*A. australiense*) | LSU, D1D2, ITS, COI |  |
|  | *Gymnodinium catenatum* | LSU, D1D2, ITS, COI |  |

|  |  |  |
| --- | --- | --- |
| *Noctiluca scintillans* | LSU, D1D2 |  |
| *Tripes (Ceratium) furca* | LSU, SSU, D1D2 | rbcl: BOLD; |
| *Tripes (Ceratium) fusus* | LSU, SSU, D1D2, COI | COI: BOLD or NCBI |
| *Tripes* sp. (genus) | SSU |  |
| *Tripes muelleri* | LSU, SSU |  |
| **Diatoms** |  |  |
| *Pseudo-nitzschia* sp. (genus) | LSU, D1D2, SSU, ITS |  |
| *Pseudo-nitzschia cuspidata* | LSU, D1D2, ITS, rbcl |  |
| *Pseudo-nitzschia pungens* | LSU, D1D2, ITS, rbcl |  |
| **Haptophytes** |  |  |
| *Emiliania huxleyi* | LSU, D1D2, rbcl, COI |  |
| **Cnidarians** |  |  |
| *Aurelia* spp. | LSU, D1D2, ITS, COI |  |
| *Cyanea* spp. | LSU, ITS, COI |  |
| *Physalia* | LSU, ITS, COI |  |
| **Molluscs** |  |  |
| *Crassostrea gigas* | LSU, D1D2, ITS, COI |  |
| *Ostrea angasi* | LSU, COI |  |
| *Mytilus galloprovincialis* | LSU, D1D2, ITS, COI |  |
| *Modiolus* spp. | LSU, D1D2, ITS, COI |  |
| **Genes involved in toxin production** |  |  |
| *SxtA* |  |  |

### 2.4 Library preparations

We prepared metagenomic libraries from all DNA extracts following Weyrich et al. (2017), with the following modifications. A 20 μL aliquot of DNA was repaired (15 min, 25 °C) in a 40 μL reaction using T4 DNA polymerase (New England Biolabs). After purifying the DNA (MinElute™ Reaction Cleanup Kit, Qiagen), a ligation step followed (T4 DNA ligase, Fermentas) in which truncated Illumina-adapter sequences containing two unique 5 base-pair (bp) barcodes were attached to the double-stranded DNA (60 min, 22 °C) (Meyer and Kircher, 2010). DNA purification (MinElute™ Reaction Cleanup Kit, Qiagen) was performed, followed by a fill-in reaction with adapter sequences (Bst DNA polymerase, New England Biolabs; 30 min, 37 °C, with polymerase deactivation for 10 min, 80 °C). For metagenomic shotgun library preparations we followed the protocol outlined in detail in Armbrecht et al. (2020), with slight modifications described in Supplementary Material Methods. For sequencing library preparations for the hybridisation capture we followed the MyBaits® Manual v4.1 April, 2018; Arbor Biosciences, USA, with modifications detailed in Supplementary Material Methods. Sequencing was performed at the Australian Cancer Research Foundation Cancer Genomics Facility & Centre for Cancer Biology, Adelaide, Australia, and at the Garvan Institute of Medical Research, KCCG Sequencing Laboratory (Kinghorn Centre for Clinical Genomics), Darlinghurst, Australia.

### 2.5 Data analysis

#### 2.5.1 Bioinformatics

Bioinformatic processing and filtering of the sequencing data, hereafter referred to as datasets ‘Shotgun’, ‘Phytobaits1’ and ‘HABbaits1’, followed established protocols previously described in Armbrecht et al. (2020), with the exception that we used the NCBI Nucleotide database (ftp://ftp.ncbi.nlm.nih.gov/blast/db/FASTA/nt.gz, downloaded November 2019) as the reference database to align our *sed*aDNA sequences to (allowing us to run all three datasets against the same database; see Supplementary Material Methods). All species detected in EBCs (Supplementary Material Table 1) were subtracted from the sample data, and hereafter the term ‘samples’ refers to sediment-derived data post-EBC subtraction. For each dataset (Shotgun, Phytobaits1 and HABbaits1), we used MEGAN6 Community Edition V6.18.10 to rank our assigned reads by domain and exported these read counts. We determined relative abundances per domain per sample, and the average and standard deviation per domain across all samples from MCS3 and GC2 (separately for each site due to relatively high variability in relative abundance between them, see results). To quantify the increase in the proportion of our target domain Eukaryota using Phytobaits1 and HABbaits1 relative to Shotgun, we determined the ratio between the average relative abundance per domain between Phytobaits1:Shotgun, and HABbaits1:Shotgun.

#### 2.5.2 Ancient DNA authenticity assessment and damage analysis

To assess the authenticity of our Shotgun, Phytobaits1 and HABbaits1 *sed*aDNA we ran the ‘MALTExtract’ and ‘Postprocessing’ tools of the HOPS v0.33-2 pipeline (Hübler et al., 2020). For specific configuration settings see Supplementary Material Methods. We processed each dataset using the ‘def_anc’ mode, which provided results for all filtered reads (‘default’; D) as well as all reads that had at least one damage lesion in their first 5 bases from either the 5’ or 3’ end (‘ancient’; A) (Hübler et al., 2020). Generally, HOPS determines DNA damage patterns separately for individual taxa, i.e., requires an input list of target taxa for which to compare the *sed*aDNA sequences identified in our samples to their modern references. We used two taxa screening lists with the aim to generate *sed*aDNA damage profiles for a representative regional phytoplankton species: (*a*) ‘Eukaryota’, to allow a general assessment of the amount of eukaryote sequences categorised as ‘default’ or ‘ancient’ in each of our samples; and (*b*) a set of selected marine organisms known to be common in our Tasmanian study region (Table 1). Subsequently to (*a*) we used the HOPS-generated ‘RunSummary’ output to determine the ratio of ancient to default in each sample for each dataset (‘A:D’ ratio hereafter). Eukaryote taxa recovered in the EBCs (Supplementary Material Table 1) were excluded from the calculation. Subsequent to (*b*), we used the MaltExtract Interactive Plotting Application (MEx-IPA, by J. Fellows Yates; https://github.com/jfy133/MEx-IPA) to visualise *sed*aDNA damage profiles of the target phytoplankton *Emiliania huxleyi* Shotgun, Phytobaits1 and HABbaits1 (ancient reads only).

#### 2.5.3 Statistics

To determine relationships between the A:D ratio and subseafloor depth and test the A:D ratio’s validity as *sed*aDNA authenticity proxy, we performed two-tailed Pearson correlation analyses between the A:D ratios of Shotgun, Phytobaits1 and HABbaits1 (n = 27 each, as no data was retrieved for 3 samples, see section 3.1) and subseafloor depth using the software PAST (Hammer et al., 2001).

## 3 Results

### 3.1 Proportions of Eukaryota in Shotgun, Phytobaits1 and HABbaits1

After filtering, we retained between 4.6 (GC2A 15 - 16.5 cm) and 16.2 M (GC2B 5 - 6.5 cm) reads per sample for Shotgun, between 0.1 (MCS3 4 - 5.5 cm) and 4.6 M (GC2A 115 - 116.5 cm) reads per sample for Phytobaits1 and between 0.2 (GC2A 45 - 46.5 cm) and 2.8 M (GC2A 115 - 116.5 cm) reads for HABbaits1. We retrieved no data for 3 out of 30 samples and these samples were excluded from downstream processing. The 3 samples with no data were MCS3 0 - 1.5 cm with Shotgun, GC2B 115 - 116.5 cm with Phytobaits1 and HABbaits1, and GC2A 85 - 86.5 cm with HABbaits1 - likely due to low template DNA concentrations. Our EBCs for Shotgun, Phytobaits1 and HABbaits1 detected a total of 121, 69 and 28 eukaryote taxa (Supplementary Material Table 1), emphasising the importance of sequencing controls and filtering *sed*aDNA data accordingly to remove contaminants.

Based on alignments using the NCBI Nucleotide database, the majority of Shotgun reads were assigned to Bacteria (86 + 5% and 63 + 16 % for MCS3 and GC2, respectively; Fig. 3a,b), and a relatively small portion to Eukaryota (5 + 2% and 28 + 15% to for MCS3 and GC2, respectively, Fig. 3a,b). This small proportion of Eukaryota increased to 21 and 53% in MCS3 and GC2 using Phytobaits1 (4.4x and 1.9x over Shotgun, respectively), and 47 and 76% in MCS3 and GC2 using HABbaits1 (9.6x and 2.7x over Shotgun respectively) (Fig. 3). Phytobaits1 and HABbaits1 were efficient in the targeted enrichment of Eukaryota *sed*aDNA from marine sediments, with comparatively little ‘bycatch’ of Bacteria and Archaea (i.e., a decrease in the proportion of Bacteria and a <2.1x increase in Archaea relative to Shotgun; Fig. 3c,d). Phytobaits1 included three cyanobacterial targets, therefore, some capture of bacterial sequences was expected; less than Shotgun but more than HABbaits1 (Fig. 3a,b).

**Figure 3:**
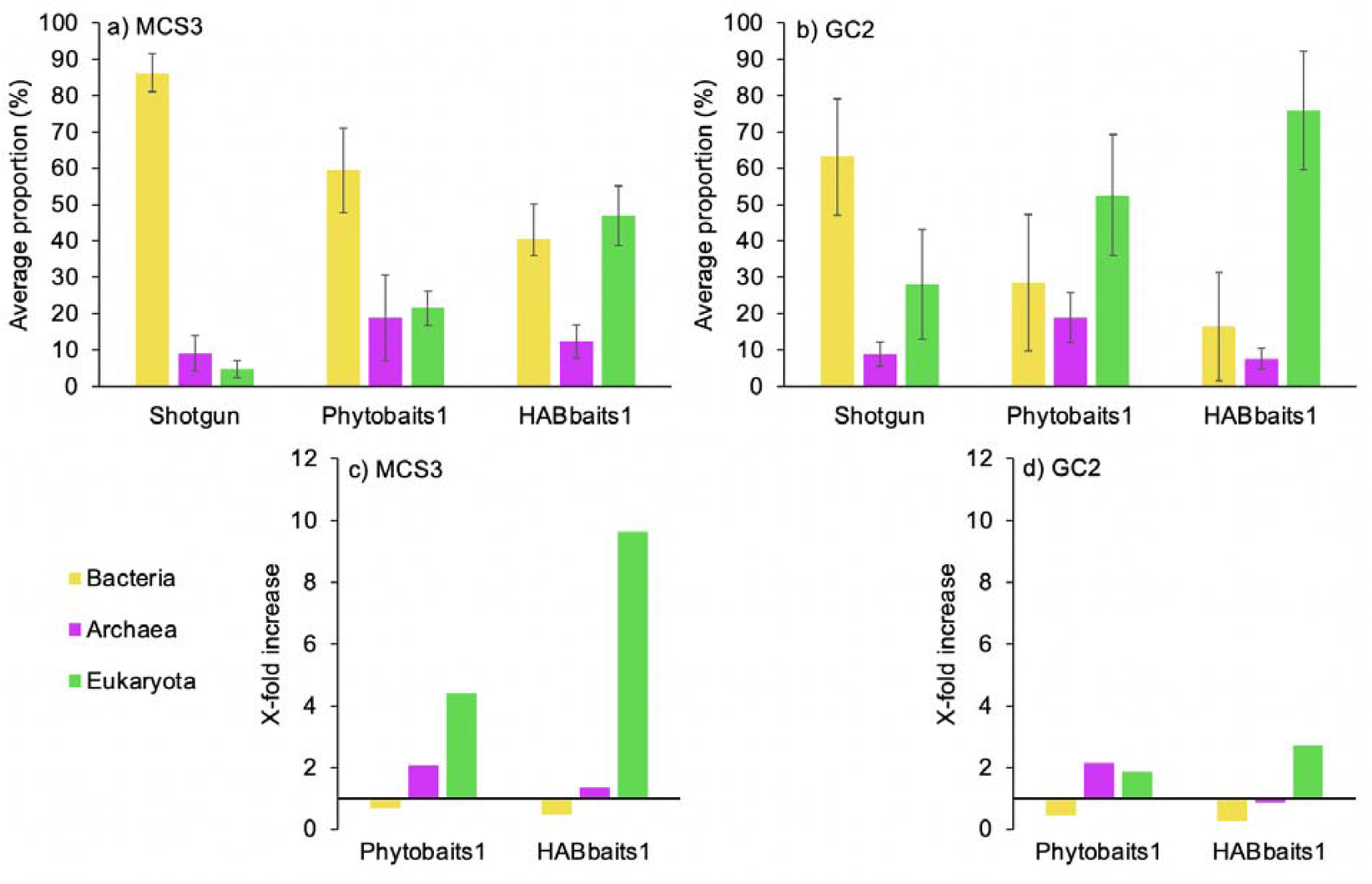
Proportions of reads assigned to Bacteria, Archaea and Eukaryota using Shotgun, Phytobaits1 and HABbaits1. a,b) Average proportion of reads and standard deviation across inshore MCS3 (n = 9) and offshore GC2 (n = 18) samples, respectively. c,d) Increase in the proportion of Bacteria, Archaea and Eukaryota in Phytobaits1 and HABbaits1 relative to Shotgun for MCS3 and GC2 samples, respectively, based on average proportions shown in (a,b).

### 3.2 Assessment of sedaDNA authenticity

For both inshore MCS3 and offshore GC2, the A:D ratio determined per sample increased with sub-seafloor depth for each of the three datasets Shotgun, Phytobaits1 and HABbaits1 (Fig. 4). At the seafloor surface, we determined an A:D ratio of approximately 0.05 for MCS3 and GC2, which slightly increased with depth until ~25 cmbsf, before a steeper increase between ~25 - 35 cmbsf, and, in offshore GC2, remained relatively stable at ~0.3 below 35 cmbsf (>1000 years of age). Correlation analyses showed that this increase of the A:D ratio with increasing subseafloor depth was highly significant for each dataset (Table 2). Additionally, the A:D ratios of the three different datasets (Shotgun, Phytobaits1 and HABbaits1) were significantly positively correlated with each other, indicating that the original proportions of *sed*aDNA damage patterns preserved in Shotgun were maintained in our hybridisation capture approach using both Phytobaits1 and HABbaits1.

**Figure 4:**
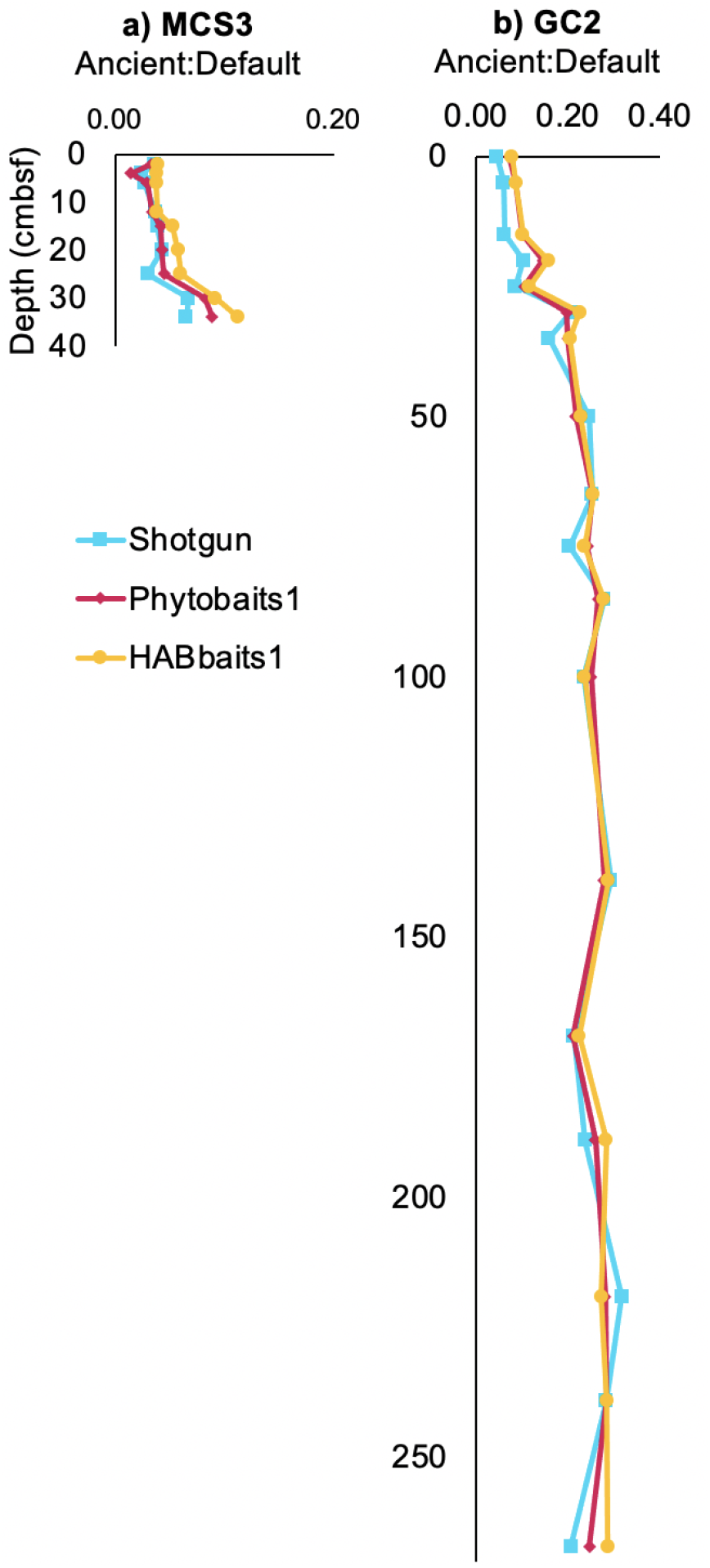
A:D ratio of reads assigned to Eukaryota with subseafloor depth. Shown is the increase in the A:D ratio with depth (centimetres below seafloor, cmbsf) in both a) MCS3, b) GC2. See Table 1 for correlation between A:D ratio per dataset (Shotgun, Phytobaits1, HABbaits1) and depth, and amongst the datasets.

**Table 2:**
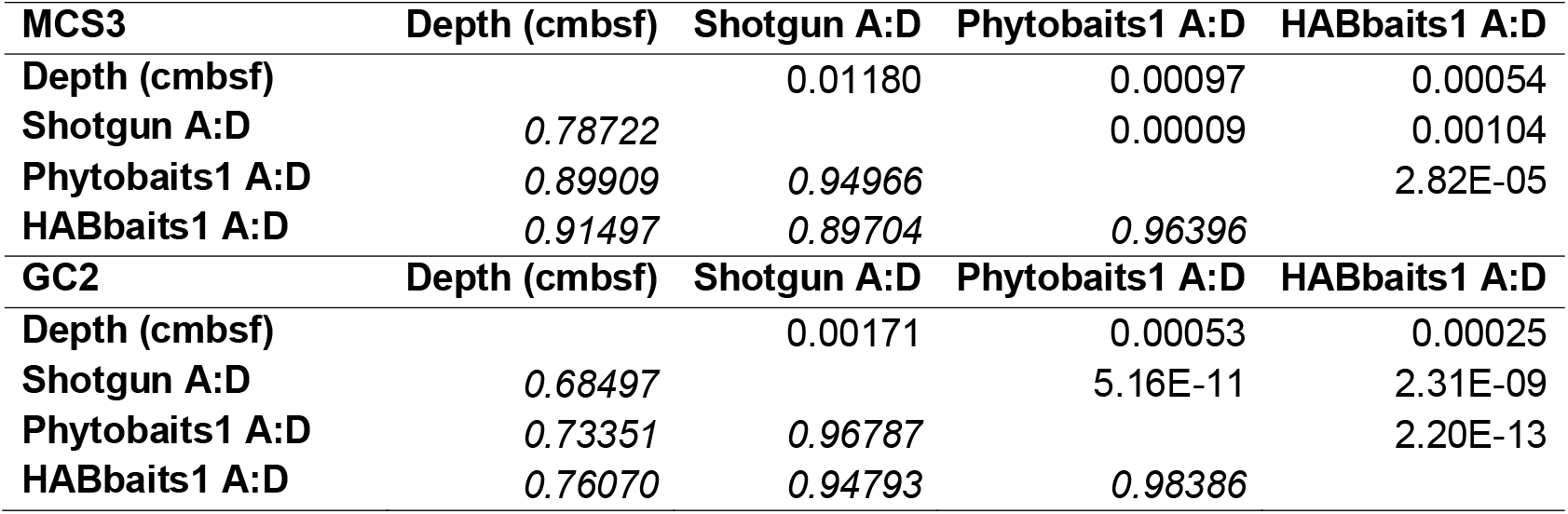
Summary statistics of correlation analysis between the A:D ratio and subseafloor depth. Pearson correlation coefficients *r* (in italics, lower matrix triangle) and corresponding two-tailed probability that *r* is uncorrelated (upper triangle of matrix; *i.e.,* all values <0.005 denote a significant correlation) between subseafloor depth (cmbsf) and Shotgun, Phytobaits1 and HABbaits1 A:D ratios (n = 27 each).

| **MCS3** | **Depth (cmbsf)** | **Shotgun A:D** | **Phytobaits1 A:D** | **HABbaits1 A:D** |
| --- | --- | --- | --- | --- |
| **Depth (cmbsf)** |  | 0.01180 | 0.00097 | 0.00054 |
| **Shotgun A:D** | *0.78722* |  | 0.00009 | 0.00104 |
| **Phytobaits1 A:D** | *0.89909* | *0.94966* |  | 2.82E-05 |
| **HABbaits1 A:D** | *0.91497* | *0.89704* | *0.96396* |  |
| **GC2** | **Depth (cmbsf)** | **Shotgun A:D** | **Phytobaits1 A:D** | **HABbaits1 A:D** |
| **Depth (cmbsf)** |  | 0.00171 | 0.00053 | 0.00025 |
| **Shotgun A:D** | *0.68497* |  | 5.16E-11 | 2.31E-09 |
| **Phytobaits1 A:D** | *0.73351* | *0.96787* |  | 2.20E-13 |
| **HABbaits1 A:D** | *0.76070* | *0.94793* | *0.98386* |  |

### 3.3 DNA damage profiles of the marine coccolithophore Emiliania huxleyi

The *sed*aDNA damage analysis provided DNA damage profiles for most of the target taxa on our selected taxa list (taxa list ‘*b*’). the number of ancient sequences assigned to the ubiquitous coccolithophore *Emiliania huxleyi* was much higher, allowing the generation of more detailed DNA damage profiles. Ancient *E. huxleyi* sequences ranged from a total of 0 - 34 reads in inshore MCS3 and 5 - 2,651 in offshore GC2 for Shotgun, from 0 - 7 in MCS3 and 1 - 947 in GC2 for Phytobaits1, and from 0 - 7 in MCS3 and 1 - 1183 in GC2 for HABbaits1. A lower representation of ‘ancient’ sequences in inshore MCS3 is consistent with our observation of a lower A:D ratio in sediments above ~35 cmbsf (i.e., the complete length of MCS3) (see section 3.2). Damage profiles for *E. huxleyi sed*aDNA are much more variable in inshore MCS3 (and in the upper ~25 cmbsf of GC2; Fig. 5,6) than the profiles of deeper, more stable offshore GC2 samples, likely resulting from a scarcity of reads in the upper sediment layers and DNA damage patterns not being as pronounced as in deeper GC2 samples.

**Figure 5:**
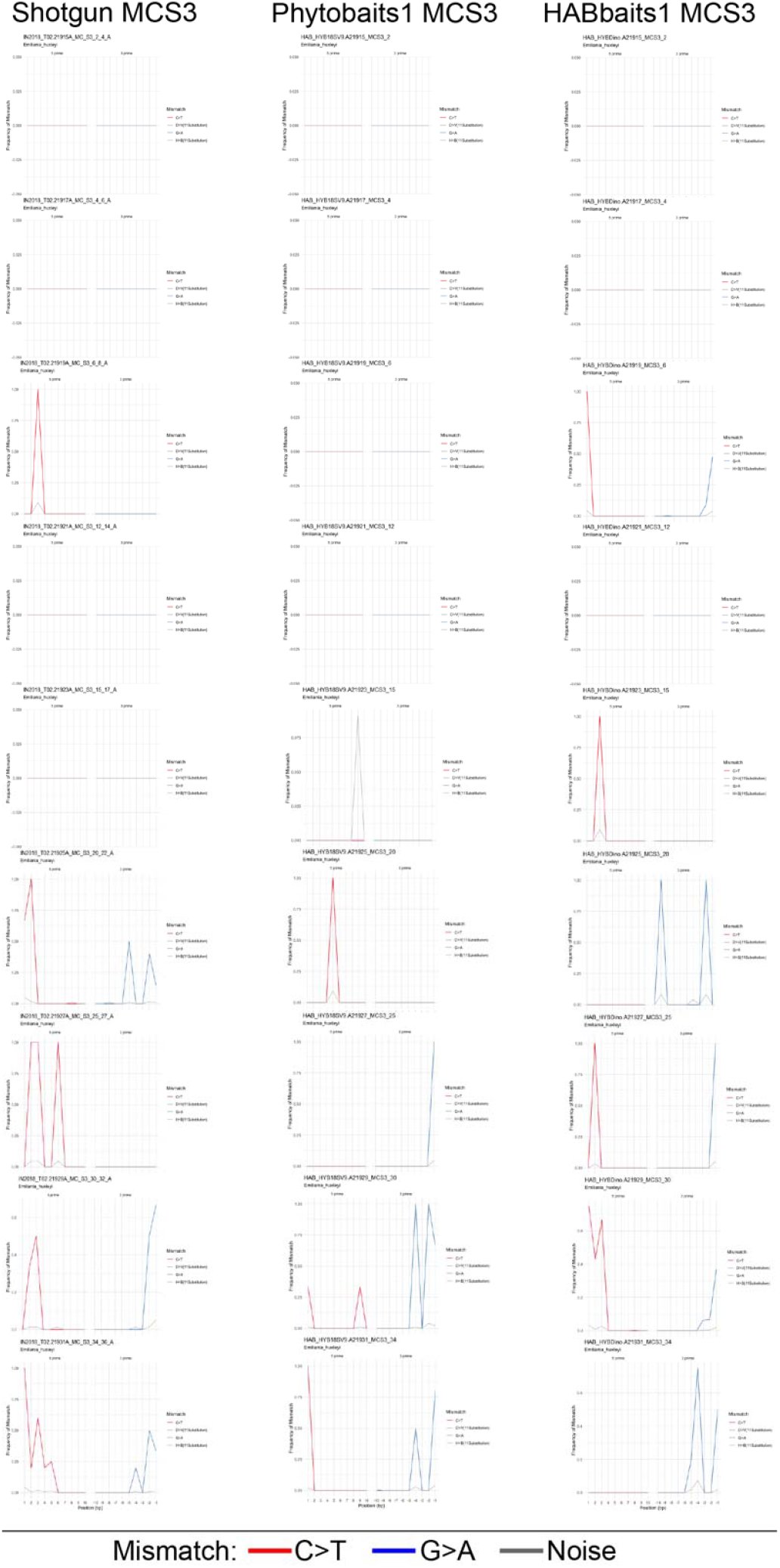
*Sed*aDNA damage profiles of *Emiliania huxleyi* in MCS3. *E. huxleyi sed*aDNA damage profiles (frequency of mismatch against base pair position) per sample for Shotgun, Phytobaits1 and HABbaits1 in MCS3 (listed from top-down). The red and blue lines denote C>T substitutions in 5’ direction and G>A substitutions in 3’ direction, respectively, for all ancient alignments. Grey lines denote estimated noise (Hübler et al., 2020).

**Figure 6:**
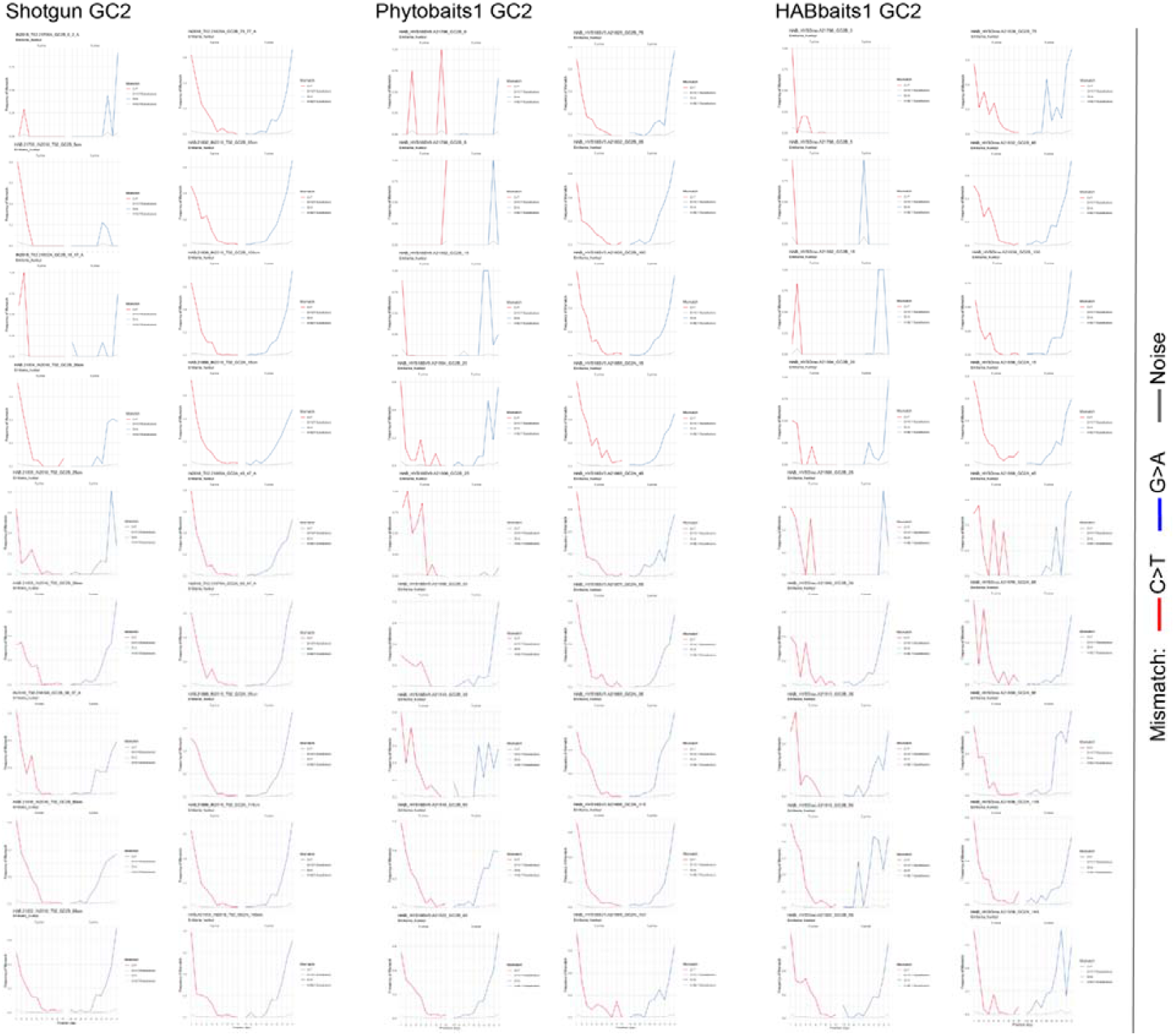
*Sed*aDNA damage profiles of *Emiliania huxleyi* in GC2. *E. huxleyi seda*DNA damage profiles (frequency of mismatch against base pair position) per sample for Shotgun, Phytobaits1 and HABbaits1 in GC2 (listed from top-down with GC2 profiles continuing in the second column for each dataset). The red and blue lines denote C>T substitutions in 5’ direction and G>A substitutions in 3’ direction, respectively, for all ancient alignments. Grey lines denote estimated noise (Hübler et al., 2020).

The *E. huxleyi sed*aDNA damage profiles of Shotgun, Phytobaits1 and HABbaits1 were consistent across samples (Fig. 5,6), suggesting that the hybridisation capture technique reliably preserves the DNA damage patterns of the original sample (represented by Shotgun) and is well-suited for the capture of past marine eukaryote *sed*aDNA. Further, HOPS provided a valid approach to authenticate *sed*aDNA from marine eukaryotes. We were unable to generate clear DNA damage profiles from the upper ~25 – 35 cmbsf in both MCS3 (spanning the last ~125 years; Fig. 5) and GC2 (~900 years, Fig. 6), indicating that DNA damage is not as pronounced in the upper (younger) sediment layers at our study location and detectable only below that depth. Below ~35 cmbsf in GC2 the *E. huxleyi* DNA damage profiles assumed a typical U-shape as the number of mismatches at the end of DNA fragments increases (Fig. 6). Our *E. huxleyi sed*aDNA damage profiles are the first generated for a marine eukaryote - and extend over an 8,000-year timescale.

## 4 Discussion

In this study we designed two new RNA bait sets and applied the hybridisation capture technique to inshore and offshore marine sediments to investigate marine eukaryotes more broadly (Phytobaits1) and in a more tailored approach focusing on selected taxa common and often harmful in our study region (HABbaits1). Our results showed that hybridisation capture improved the genetic yield of eukaryote *sed*aDNA, and preserved DNA damage patterns that allowed us to make an assessment of *sed*aDNA authenticity, as well as enabling us to generate the first ancient DNA damage profiles of a keystone marine phytoplankton organism, the ubiquitous coccolithophore *Emiliania huxleyi.*

### 4.1 Targeted enrichment of marine Eukaryota sedaDNA

Both Phytobaits1 and HABbaits1 successfully captured *sed*aDNA of eukaryote organisms in two sediment cores collected off the Tasmanian east coast. Eukaryote *sed*aDNA has been shown repeatedly to be present in low amounts in seafloor sediments, which has limited the metagenomic analysis and detailed reconstruction of past marine ecosystems. While both Phytobaits1 and HABbaits1 achieved a considerable enrichment in Eukaryota for our inshore site MCS3 (4- to 9-fold, respectively), this increase was about half at the offshore site GC2. The difference in Eukaryota increase may be due to the initial difference in Eukaryota proportions at the two sites. Shotgun showed Eukaryota contributed ~5% to the total pool of *sed*aDNA at MCS3, while contributing ~28% at GC2. The latter high proportion is primarily a result of a sharp increase in the relative abundance of Eukaryota in GC2 below 35 cmbsf. It is possible that this initially relatively high presence of Eukaryota sequences in the GC2 *sed*aDNA extracts saturated the baits in our hybridisation reaction. This would explain a less pronounced increase in GC2 Eukaryota proportions using either bait set. To further increase the Eukaryota signal in future studies, it may be beneficial to add a larger volume of baits (>3 μL) to *sed*aDNA extracts that either are expected or have been shown (e.g., by shallow shotgun sequencing prior to enriching) to have a relatively high Eukaryota *sed*aDNA content.

### 4.2 Assessment of sedaDNA authenticity Interesting

The HOPS bioinformatic tool (Hübler et a., 2020) proved highly valuable in identifying and analysing ancient eukaryote sequences in our *sed*aDNA. The HOPS generated output of our ‘Eukaryota’ (taxa list *a*) run enabled the determination of the A:D ratio, a parameter that can be used as a proxy of *sed*aDNA authenticity in the future. Here, the A:D ratio was quite low (0.05 – 0.3), which might point towards our *sed*aDNA being relatively well preserved, and thus a high proportion of reads passing the default filtering criteria. The latter criteria used a minimum percent identity (mpi) level of 95%, a relatively stringent cut-off while still retaining the majority of reads. Increasing the mpi cut-off may have resulted in a higher A:D ratio due to less reads passing the filtering criteria, however, this would also eliminate the majority of reads available for analyses.

At both our inshore and offshore site, we observed a significant increase in A:D ratio with subseafloor depth, demonstrating that eukaryote *sed*aDNA shows increased DNA damage with increasing age of sediments. However, at both sites the A:D ratio was consistently lower in the upper 25 - 35 cmbsf (~0.05), then increased sharply down-core from this depth in both (~0.3), and remained at this level towards the bottom of in GC2. Whether reaching such a ‘limit’ in the A:D ratio is a pattern characteristic of our study location, or indicative of a ‘critical depth’ below which *sed*aDNA degradation accelerates, remains to be investigated. Future *sed*aDNA studies should investigate how the A:D ratio varies in much older sediment records (older than Holocene) and depending on sediment properties (e.g., clay-rich sediments that appear to benefit DNA preservation; Vuillemin et al., 2019).

The strong positive correlation between the A:D ratios amongst Shotgun, Phytobaits1 and HABbaits1, demonstrates that DNA damage signals present in *sed*aDNA are preserved throughout the hybridisation capture approach. This is important as it allows the authentication of *sed*aDNA using bioinformatic tools (see section 4.3), which any ancient DNA study should incorporate (Hübler et al., 2020). Through hybridisation capture more target sequences are available as input for DNA damage analysis software such as HOPS, which increases the robustness of such analyses (Hübler et al., 2020), thus is strongly recommended for *sed*aDNA analyses. While future refinement of A:D ratios may be necessary, our analyses show that it can be used as a proxy for *sed*aDNA authenticity in sediment records. Generally, for marine *sed*aDNA investigations of eukaryote taxa, the capacity to assess DNA damage provides a crucial advantage over metabarcoding where DNA damage-based authenticity assessment is impossible.

### 4.3 Emiliania huxleyi sedaDNA damage profiles

Running our data through the HOPS pipeline (taxa list *b*) and MEx-IPA allowed us to generate DNA damage plots for a key marine phytoplankton species, *Emiliania huxleyi.* This ubiquitous calcareous nanoplankton has thrived in the oceans since the Cretaceous, is one of the most abundant phytoplankton species in the global ocean and is ubiquitous from tropical to temperate to Antarctic Australian waters (Hallegraeff 1984; Cubillos et al. 2007). Consistent with its biogeographic distribution in the modern ocean, we expected to detect traces of this species in our *sed*aDNA, and in higher relative abundances offshore.

We retained the maximum number of reads throughout our analyses (by examining proportions rather than rarefying our data), which enabled us to generate *E. huxleyi* DNA damage profiles from all three datasets, Shotgun, Phytobaits1 and HABbaits1. The damage profiles generated by Shotgun, Phytobaits1 and HABbaits1 per sample were very similar, indicating preservation of DNA damage patterns in our original sample (Shotgun) and in our enriched samples after hybridisation capture. Consistent with our finding of low A:D ratio in the upper 25 - 30 cmbsf, no clear *E. huxleyi* damage patterns could be determined from these depths. *sed*aDNA damage patterns with a typical U-shape were found only below ~25 cmbsf in GC2, again suggesting the existence of a critical depth below which DNA degradation becomes more pronounced, reinforcing the importance of investigating whether this phenomenon is of wider importance, and possibly correlated with the age or physical or chemical properties of marine sediments.

### 4.4 Significance of hybridisation capture and marine sedaDNA damage analysis

The study of marine *sed*aDNA offers huge potential for the comprehensive reconstruction of past marine ecosystems (including viruses, archaea, prokaryotes and eukaryotes). Eukaryotes (phytoplankton and higher organisms) are particularly popular study organisms due to their importance as primary producers and use as environmental indicators. However, *sed*aDNA studies focussing on eukaryotes have been severely limited by the very low amounts of DNA preserved in the subseafloor, and the lack of bioinformatic tools to authenticate these miniscule amounts of eukaryote *sed*aDNA in metagenomic data. To date, no marine *sed*aDNA study exists that had proven authenticity (i.e., the DNA recovered is ancient and free from modern contamination) through bioinformatic approaches such as *sed*aDNA damage analysis, a routine procedure in ancient DNA studies focussing on humans and megafauna. Our study provides a key advance in that we (*1*) used a hybridisation capture technique to enrich target marine eukaryote *sed*aDNA independent of DNA fragment size, and (*2*) applied the recently developed bioinformatic tool HOPS for *sed*aDNA damage analysis and to authenticate our marine *sed*aDNA. These advances are of importance as we are now able to bioinformatically discriminate the authentic *sed*aDNA component to more accurately estimate paleo-community composition.

## Conclusions

In this study we show the reliability of the hybridisation capture as a novel tool for investigating changing patterns of abundance of marine eukaryotes from their *sed*aDNA in seafloor sediments. We furthermore applied a new bioinformatic approach for metagenomic *sed*aDNA damage analysis, which allowed us to develop a new proxy for *sed*aDNA authenticity (the A:D ratio) that changes with subseafloor depth. Through our *sed*aDNA damage analysis were also able to generate *sed*aDNA damage profiles of the ubiquitous coccolithophore *E. huxleyi,* the first ever such profiles generated for a marine eukaryote – extending over an 8,000-year timescale. Our study provides a major step forward for the future investigation of eukaryotes from marine *sed*aDNA, enabling detailed insights into past marine ecosystem composition over geological timescales.

## Acknowledgements

We are grateful to Brian Brunelle from Arbor Biosciences, USA, for his expert assistance with designing Phytobaits1 and HABbaits1. We thank Oscar Estrada-Santamaria, Steve Richards, Holly Heiniger, Nicole Moore and Steve Johnson for their help and advice during extractions, library preparations, and hybridisation capture. We are grateful to Raphael Eisenhofer, Vilma Pérez, Yassine Souilmi, Yichen Liu, and Ron Hübler for their help with on bioinformatic analyses. We thank the Marine National Facility, the crew of RV *Investigator* voyage IN2018_T02 and the scientific team for their support during field work (2018 MNF Grant H0025318), and Prof. Andrew McMinn for assistance with collection of the cores. This study was funded through the 2017 Australian Research Council (ARC) Discovery Project DP170102261. AC was funded by ARC Laureate Fellowship FL140100260.

## Author contributions

L.A. designed this research, carried out laboratory work, bioinformatic and statistical analyses and wrote the first draft of the manuscript. G.H., C.B., and C.W. collected and provided the core samples. C.W. provided sediment core dating. A.C. provided guidance on the hybridisation capture technique and ancient DNA analyses. G.H., C.B. and A.C. assisted with data interpretation and secured funding for this project. All co-authors provided comments and feedback on manuscript drafts, and edited the final manuscript submitted for publication.

## Competing interests

All co-authors declare that there are no competing interests.

## Materials & Correspondence

Linda Armbrecht

## Notes

### Competing Interest Statement

The authors have declared no competing interest.

## References

Armbrecht, L.H., Coolen, M.J., Lejzerowicz, F., George, S.C., Negandhi, K., Suzuki, Y., Young, J., Foster, N.R., Armand, L.K., Cooper, A. and Ostrowski, M., 2019. Ancient DNA from marine sediments: Precautions and considerations for seafloor coring, sample handling and data generation. Earth-Science Reviews 196, 102887.

Armbrecht, L., Herrando-Pérez, S., Eisenhofer, R., Hallegraeff, G.M., Bolch, C.J. and Cooper, A., 2020. An optimized method for the extraction of ancient eukaryote DNA from marine sediments. Molecular Ecology Resources 20, 906–919.

Armbrecht, L. 2020. The Potential of sedimentary ancient DNA to reconstruct past ocean ecosystems. Oceanography, https://doi.org/10.5670/oceanog.2020.211.

Collin, T.C., Drosou, K., O’Riordan, J.D., Meshveliani, T., Pinhasi, R. and Feeney, R.N.M., 2020. An open-sourced bioinformatic pipeline for the processing of Next-Generation Sequencing derived nucleotide reads: Identification and authentication of ancient metagenomic DNA. BioRxiv, https://doi.org/10.1101/2020.04.20.050369.

Cubillos, J. C., Wright, S. W., Nash, G., de Salas, M. F., Griffiths, B., Tilbrook, B., Poisson, A. & Hallegraeff, G. M. 2007. Calcification morphotypes of the coccolithophorid *Emiliania huxleyi* in the Southern Ocean: changes in 2001 to 2006 compared to historical data. Marine Ecology Progress Series 348, 47–54.

De Schepper, S., Ray, J.L., Skaar, K.S., Sadatzki, H., Ijaz, U.Z., Stein, R. and Larsen, A., 2019. The potential of sedimentary ancient DNA for reconstructing past sea ice evolution. The ISME Journal 13, 2566–2577.

De Vargas, C., Audic, S., Henry, N., Decelle, J., Mahé, F., Logares, R., Lara, E., Berney, C., Le Bescot, N., Probert, I. and Carmichael, M., 2015. Eukaryotic plankton diversity in the sunlit ocean. Science 348, 1261605, and companion website http://taraoceans.sb-roscoff.fr/EukDiv/.

Foster, N.R., Gillanders, B.M., Jones, A.R., Young, J.M. and Waycott, M., 2020. A muddy time capsule: using sediment environmental DNA for the long-term monitoring of coastal vegetated ecosystems. Marine and Freshwater Research 71, 869–876.

Ginolhac, A., Rasmussen, M., Gilbert, M.T.P., Willerslev, E. and Orlando, L., 2011. mapDamage: testing for damage patterns in ancient DNA sequences. Bioinformatics 27, 2153–2155.

Guillou, L., Bachar, D., Audic, S., Bass, D., Berney, C., Bittner, L., Boutte, C., Burgaud, G., de Vargas, C., Decelle, J. and del Campo, J., 2012. The Protist Ribosomal Reference database (PR^2^): a catalog of unicellular eukaryote small sub-unit rRNA sequences with curated taxonomy. Nucleic Acids Research 41, D597–D604.

Grzebyk, Daniel; Audic, Stéphane; Decelle, Johan; de Vargas, Colomban (2017), “PHYTOPK28-D1D2: A curated database of 28S rRNA gene D1-D2 domains from eukaryotic organisms dedicated to metabarcoding analyses of marine phytoplankton samples”, Mendeley Data v1 http://dx.doi.org/10.17632/mndb4h87yg.1

Hallegraeff, G.M. 1984 Coccolithophorids (calcareous nanoplankton) from Australian waters, Botanica Marina 27, 229–247.

Hammer, Ø., Harper, D.A.T., and P. D. Ryan, 2001. PAST: Paleontological Statistics Software Package for Education and Data Analysis. Palaeontologia Electronica 4, 9.

Hays, G.C., Richardson, A.J. and Robinson, C., 2005. Climate change and marine plankton. Trends in Ecology and Evolution 20, 337–344.

Horn, S. 2012. Target enrichment via DNA hybridisation capture. In: Ancient DNA, Methods and Protocols, eds. B. Shapiro and M. Hofreiter, Springer New York, Dordrecht, Heidelberg, London, pp. 177–188.

Hübler, R., Key, F.M., Warinner, C., Bos, K.I., Krause, J. and Herbig, A., 2019. HOPS: Automated detection and authentication of pathogen DNA in archaeological remains. Genome Biology 20, 1–13.

Jónsson, H., Ginolhac, A., Schubert, M., Johnson, P.L. and Orlando, L., 2013. mapDamage2. 0: fast approximate Bayesian estimates of ancient DNA damage parameters. Bioinformatics 29, 1682–1684.

Murchie, T.J., Kuch, M., Duggan, A.T., Ledger, M.L., Roche, K., Klunk, J., Karpinski, E., Hackenberger, D., Sadoway, T., MacPhee, R. and Froese, D., 2020. Optimizing extraction and targeted capture of ancient environmental DNA for reconstructing past environments using the PalaeoChip Arctic-1.0 bait-set. Quaternary Research, https://doi.org/10.1017/qua.2020.59.

Li, C., Corrigan, S., Yang, L., Straube, N., Harris, M., Hofreiter, M., White, W.T. and Naylor, G.J., 2015. DNA capture reveals transoceanic gene flow in endangered river sharks. Proceedings of the National Academy of Sciences 112, 13302–13307

Llamas, B., P. Brotherton, K.J. Mitchell, J.E. Templeton, V.A. Thomson, J.L. Metcalf, K.N. Armstrong, M. Kasper, S.M. Richards, A.B. Camens, and M.S. Lee. 2015. Late Pleistocene Australian marsupial DNA clarifies the affinities of extinct megafaunal kangaroos and wallabies. Molecular Biology and Evolution 32, 74–584.

More, K.D., W.D. Orsi, V. Galy, L. Giosan, L. He, K. Grice, and M.J. Coolen. 2018. A 43 kyr record of protist communities and their response to oxygen minimum zone variability in the Northeastern Arabian Sea. Earth and Planetary Science Letters 496, 248–256.

MyBaits® Manual v.4.01 — Hybridization Capture for Targeted NGS, 2018. https://arborbiosci.com/wp-content/uploads/2019/08/myBaits-Manual-v4.pdf

Paijmans, J.L., Gilbert, M.T.P. and Hofreiter, M., 2013. Mitogenomic analyses from ancient DNA. Molecular Phylogenetics and Evolution 69, 404–416.

Pedersen, M.W., Overballe-Petersen, S., Ermini, L., Sarkissian, C.D., Haile, J., Hellstrom, M., Spens, J., Thomsen, P.F., Bohmann, K., Cappellini, E. and Schnell, I.B., 2015. Ancient and modern environmental DNA. Philosophical Transactions of the Royal Society B: Biological Sciences 370, 20130383.

Ratnasingham, S. and Hebert, P.D., 2007. BOLD: The Barcode of Life Data System (http://www.barcodinglife.org). Molecular Ecology Notes 7, 355–364.

Szczuciński, W., Pawłowska, J., Lejzerowicz, F., Nishimura, Y., Kokociński, M., Majewski, W., Nakamura, Y. and Pawlowski, J., 2016. Ancient sedimentary DNA reveals past tsunami deposits. Marine Geology 381, 29–33.

Taberlet, P., Coissac, E., Pompanon, F., Brochmann, C. and Willerslev, E., 2012. Towards next-generation biodiversity assessment using DNA metabarcoding. Molecular Ecology 21, 2045–2050.

Tobler, R., A. Rohrlach, J. Soubrier, P. Bover, B. Llamas, J. Tuke, N. Bean, A. Abdullah-Highfold, S. Agius, A. O’Donoghue, and I. O’Loughlin. 2017. Aboriginal mitogenomes reveal 50,000 years of regionalism in Australia. Nature 544, 180–184.

Verity, P.G. and Smetacek, V., 1996. Organism life cycles, predation, and the structure of marine pelagic ecosystems. Marine Ecology Progress Series 130, 277–293.

Vuillemin, A., Wankel, S.D., Coskun, Ö.K., Magritsch, T., Vargas, S., Estes, E.R., Spivack, A.J., Smith, D.C., Pockalny, R., Murray, R.W. and D’Hondt, S., 2019. Archaea dominate oxic subseafloor communities over multimillion-year time scales. Science Advances 5, p. eaaw4108, https://doi.org/10.1126/sciadv.aaw4108.

Weyrich, L.S., Duchene, S., Soubrier, J., Arriola, L., Llamas, B., Breen, J., Morris, A.G., Alt, K.W., Caramelli, D., Dresely, V. and Farrell, M., 2017. Neanderthal behaviour, diet, and disease inferred from ancient DNA in dental calculus. Nature 544, 57–361.

Willerslev, E. and Cooper, A., 2005. Ancient DNA. Proceedings of the Royal Society B: Biological Sciences 272, 3–16.

