## Supplementary Material for "Hybridisation capture allows DNA damage analysis of ancient marine eukaryotes"

**Contents** **Page**

| | |
|---|---|
| Methods - *SedaDNA extractions* | 2 |
| Methods - *RNA-baits design: Phytobaits1* | 3 |
| Methods - *RNA-baits design: HABbaits1* | 4 |
| Methods - *Library preparation: Metagenomic* | 5 |
| *Methods - Library preparation: Hybridisation capture* | 6 |
| Methods - *Bioinformatics: Raw sequencing data processing* | 9 |
| *Methods - Bioinformatics:* sed*aDNA authenticity assessment and damage analysis* | 10 |
| References | 11 |
| Supplementary Material Tables - Supplementary Material Table 1 | 12 |

### Methods

#### *SedaDNA extractions*

Our extraction method followed the optimised ("combined") approach outlined in detail in Armbrecht et al. (2020). In brief, we used 0.25 g of each sample and incubated in 0.75 mL EDTA overnight on a rotary mixer (room temperature, RT). After centrifuging (3 min, 13,000 rpm), we kept the supernatant (RT) while proceeding with bead-beating of the pellet (3 x 20 s with 5 s breaks) in 0.75 mL customary beat-beating and C1 solution (DNeasy PowerLyzer PowerSoil Kit, Qiagen) using a Precellys 24 homogenizer (Bertin Instruments, France) and a FastPrep FP120 (Thermo Savant, USA) for the MCS3 and GC2 samples, respectively, followed by centrifugation (3 min, 10,319 rpm). We combined purified DNA-solutions from EDTA and bead-beating at 0.75 mL each per sample, added this DNA to 6 mL modified QG binding buffer (Qiagen) with 100 µL liquid silica (Brotherton et al., 2013) in a 15 mL centrifugation tube, and stirred the solution on a rotary mixer (1 hr, RT). After centrifuging (1 min, 14,000 rpm), we resuspended and washed the pellet with 0.9 mL of QG binding buffer, and twice with 80% EtOH with each wash step followed by centrifugation (1 min, 14,000 rpm). DNA pellets were then dried (15 min, 37 °C) and resuspended in 100 µL TLE Buffer (50 µL Tris HCL (1M), 10 µL EDTA (0.5M), 5 mL nuclease-free water). Following incubation (10 min, 50 °C), we centrifuged (1 min, 14,000 rpm) and stored the supernatant (free of silica) in a sterile Lo-bind tube (Eppendorf) at -20 °C. To monitor laboratory contamination, we used extraction blank controls (EBCs) by processing 1-2 (depending on the extraction-batch size) empty bead-tubes through the extraction protocol. A total of 30 extracts were generated from sediment samples and 7 extracts from EBCs.

#### ***RNA-baits design***

*Phytobaits1*

To design Phytobaits1 we downloaded the W2_V9_PR2 database (containing 18S-V9 rDNA and rRNA sequences of marine protists and their predators, De Vargas et al., 2015; downloaded on 30 July 2018), deduplicated using Geneious software (Geneious NZ), and filtered the remaining sequences to keep only those from major phyto- and zooplankton groups (Main Text Table 1). In collaboration with Arbor Biosciences, USA, we designed RNA baits based on these 15,035 target sequences by masking any repeating Ns (i.e., any consecutive Ns that were <10 in a row were converted to Ts, with ultimately 0.1% masked), padding short targets to 84 nucleotides (nt) (i.e., any target less than 84 nt was padded with Ts up to 84 nt in length). This procedure provided 41,798 raw baits of 80 nt with 3 x tiling (creating an even coverage, i.e., one bait every ~27 nt). The raw baits were BLASTed against ArborBioscience's in-house RefSeq database containing 5,584 bacterial genome and plasmid sequences (downloaded from NCBI, May, 2018), and any baits leading to hits were removed (except for 785 loci from cyanobacterial taxa that we intended to keep, see below). This filtering step provided 36,836 baits, which were collapsed into 15,942 final baits (i.e., eliminating redundancy based on identity and overlap; using >83% overlap, and >95% identity). We added five 16S-V4 rRNA sequences (the prokaryotic equivalent of the small subunit ribosomal rRNA gene) of common marine cyanobacteria (one *Trichodesmium erythraeum* sequence, and two *Prochlorococcus marinus* and *Synechococcus* sp. sequences each), acquired from the SILVA database; Main Text Table 1). To check and ensure target-taxon specificity, these five cyanobacterial sequences were mapped against a non-target sequence (*Escherichia coli* 16S RefSeq sequence NR_114042.1), then reverse-transcribed to DNA, and BLASTed to the same NCBI RefSeq database described above. BLAST hits of <60 bp alignment length and <80% identity were removed, and only those baits with <50

BLAST hits were kept, resulting in 10 cyanobacterial baits. Consequently, Phytobaits1 contained a total of 15,952 RNA baits targeting the 18SV9 region of a broad diversity of phytoplankton and their predators and the 16SV4 region of three cyanobacteria.

*HABbaits1*

To design HABbaits1 we manually collated a total of 805 LSU, SSU, D1-D2-LSU, COI, rbcL and ITS sequences for specific marine target organisms often associated with harmful algal bloom events in our study region, primarily dinoflagellates but also certain diatoms, a coccolithophore, jelly- and shellfish and the saxitoxin A4 gene, involved in paralytic shellfish toxin production by the dinoflagellates *Gymnodinium catenatum* and some species of the genus *Alexandrium* (Main Text Table 1). As with Phytobaits1, we worked in collaboration with Arbor Biosciences, USA, to design RNA baits based on the collated sequences (converting consecutive (<10) Ns to Ts and RNA sequences to DNA, masking input sequences for simple repeats (0.4%)), attaining 23,064 raw 80 nt baits (using 3x tiling, as for Phytobaits1, see above). Each bait candidate was BLASTed against three target genomes (the oyster *Crassostrea gigas,* coccolithophore *Emiliania huxleyi,* mussel *Mytilus galloprovincialis*), and four non-target genomes (diatoms *Fragilariopsis cylindrus, Phaeodactylum tricornutum,* dinoflagellate *Symbiodinium minutum,* diatom *Thalassiosira pseudonana,* jellyfish *Clytia hemisphaerica*), and a hybridisation melting temperature ($T_m$)* was estimated for each hit assuming standard myBaits® buffers and conditions ($T_m$ is defined as the temperature at which 50% of molecules are hybridised). For each target bait candidate, one BLAST hit with the highest $T_m$ was first discarded from the results (allowing for 1 hit in the genome), and only the top 500 hits (by bit score) were considered. Based on the distribution of remaining calculated $T_m$'s, we filtered out non-specific baits using stringent (only specific baits pass) criteria (i.e., bait candidates pass if they satisfy one of these conditions: a) no hits with $T_m$

above 60°C, b) ≤2 hits 62.5 – 65°C, c) ≤10 hits 62.5 – 65°C and at least 1 failing flanking bait, d) ≤10 hits 62.5 – 65°C, 2 hits 65 – 67.5°C, and <2 passing flanking baits, e) ≤2 hits 62.5 – 65°C, 1 hit 65 – 67.5°C, 1 hit 70°C or above, and <2 passing flanking baits. Bait candidates were removed when a hit was determined after BLASTing them against the non-target genomes. This stringent filtering procedure was applied du ensure maximum target-specificity, and resulted in a total of 15,310 baits for HABbaits1.

#### ***Library preparations***

*Metagenomic shotgun sequencing library preparations*

We followed the shotgun sequencing library preparation protocol detailed in Armbrecht et al. (2020), with the following modifications. We used 3 µL of the Bst-reaction product as input for a 25 µL PCR with the primers IS7 and IS8 (Meyer and Kircher, 2010) (preparing 5 PCR replicates for most samples; however, for twelve offshore core samples (and two associated EBCs) we ran 8 PCR replicates, and for three MCS3 samples (0 - 1.5 cm, 12 - 13.5 cm, 34 - 35.5cm) and the associated EBC we only ran duplicates; with the number of PCR replicates varying due to DNA template limitations resulting from previous PCR trials). Each PCR reaction included 14.2 nuclease-free $H_2O$, 2.5 µL 10× Gold Buffer, 2.5 µL 25mM $MgCl_2$, 0.25 µL 25mM dNTPs, 1.25 IS7, 1.25 IS8, and 0.1 µL AmpliTaq Gold Polymerase (Applied Biosystems). Thermal cycling specifications were as follows: 6 min at 94 °C, 18 cycles (22 cycles in the case of the 12 GC2 samples and the two associated EBCs) of 30 s denaturation at 94 °C, 30 s annealing at 60 °C, 40 s extension at 72 °C, and 10 min of final extension. We purified our PCR products using AxyPrep magnetic beads (Axygen Biosciences; 1:1.8 library:beads), eluted the DNA in Buffer EB (Qiagen) with 0.05% Tween®20 (Sigma Aldrich), and quantified DNA concentrations using Qubit™ dsDNA HS Assays (Molecular Probes). Next, we ran additional PCRs (25 µL reactions, 5 replicates each) using the same reagents as above but with the Indexing Primer IS4, a GAII index (Meyer and Kircher, 2010) and 13 cycles. We pooled the PCR products per sample, cleaned them using AxyPrep magnetic beads (1:1.1 library:beads), and performed quantity and quality checks through TapeStation (Agilent Technologies, USA). For those samples showing primer-dimer, we repeated the AxyPrep clean-up (1:1.1 library:beads) and TapeStation control. We prepared two equimolar (7.3 nM and 10 nM as per TapeStation) sequencing pools, to which we added the libraries prepared from EBCs in a 1 in 10 dilution. We submitted the final pools for Illumina NextSeq sequencing (2 × 75 bp cycle) at the Australian Cancer Research Foundation Cancer Genomics Facility & Centre for Cancer Biology, Adelaide, Australia, and at the Garvan Institute of Medical Research, KCCG Sequencing Laboratory (Kinghorn Centre for Clinical Genomics) Darlinghurst, Australia.

***Library preparations for/and hybridisation capture***

A minimum of 100 ng DNA in 7 µL is recommended as input for hybridisation capture reactions as per the manufacturer's instructions (MyBaits® Manual v4.1 April, 2018; Arbor Biosciences, USA). Based on pilot trials with three marine sediment samples (not shown), we determined that the minimum input can be reduced to ~50 ng if DNA concentrations are very low, as was the case for our samples. To achieve at least ~50 ng input DNA in 7 µL, we re-amplified remaining sample material from most of our cleaned post-IS7/IS8 PCR products (see section shotgun library prep. above) in another IS7/IS8 PCR (one 75 µL reaction with 9 µL DNA input per sample, using the same reagent composition as described in shotgun libraries above and 10 amplification cycles). We combined the barcoded EBCs (1 µL each, using a 1 in 10 dilution of EBC_A24029 due to its comparably high DNA concentration relative to the other EBCs) in one PCR reaction (7 µL EBC DNA template total) for the downstream enrichments. After re-amplification, the PCR products were cleaned using AxyPrep magnetic beads (1:1.8 library:beads) and quantified using Qubit DNA assays. Samples for which the initial IS7/IS8 PCR provided comparatively high DNA concentrations were not re-amplified prior to hybridisation capture. Using this procedure, we generated 62.53 $\pm$ 25.92 ng of DNA (23.24 – 171.75 ng; 0.07 ng for the EBC pool) for use as input material for the hybridisation capture with Phytobaits1 and HABbaits1.

We followed the manufacturer's protocol for hybridisation capture with the following modifications. In the Hybridisation Mix ("HYBs") we used 3 µL baits per reaction, and in the Blockers Mix ("LIBs") we used the blockers Nimblegen SeqCapEZ (a plant repetitive elements blocker), Block O and Block A (Salmon Sperm DNA and P5/P7 block, respectively, both provided with the MyBaits® kit), and we added 7 µL of DNA template. We combined LIBs and HYBs per sample in a Thermocycler (Thermoscientific) once the latter had been at hybridisation temperature for 5 min. For Phytobaits1 we set the hybridisation temperature to 60 °C as per the manufacturer's recommendation for short and damaged DNA molecules, and the hybridisation reaction to 40 h. For HABbaits1, we set the hybridisation temperature to 65 °C for the first 3 h to favour highly specific binding, followed by a decrease to 60 °C for the remaining 37 h of the hybridisation capture reaction. We prepared the beads for batches of 8 reactions in 1.7 mL tubes by washing the beads twice with binding buffer, then adding binding buffer and 48 µL yeast tRNA (= 480 µg per 240 mL beads) in a third washing step, followed by brief vortexing and incubation of the solution on a rotary mixer (30 min, room temperature), pelleting on a magnetic rack, and two more washes with binding buffer. We performed bead-hybrid binding for 20 min at 60 °C, with agitation by pipette-mixing, and briefly centrifuging to collect after 5 min. Subsequent washes and library resuspensions (in 40 µL Buffer EBT (EB (Qiagen) with 0.05% Tween®20 (Sigma Aldrich)) were performed as per protocol for non-KAPA HiFi HotStart polymerase amplification (incubation at 95 °C, pelleting of beads and collection of DNA containing buffer EBT).

GaII Indexing PCRs (using different indices for HABbaits1 and Phytobaits1) were performed as for shotgun sequencing libraries above, but we used one 100 µL reaction and 16 amplification cycles per sample. Initially, we used 12 and 24 µL hybridisation capture DNA template generated from MCS3 and GC2 samples as we assumed relatively high and low DNA concentrations, respectively. For samples GC2B 15-16.5 cm, GC2B 75-76.5 cm and GC2A 65-66.5 cm we used 12 µL DNA template due to previous experimental trials. Following amplification, very low DNA concentrations were determined for all HABbaits1 samples and Phytobaits1 samples MCS3 2 - 3.5 cm, MCS3 4 - 5.5cm, GC2B 85 - 86.5 cm and GC2A 75 - 76.5 cm. Therefore, we used the remaining hybridisation capture material (26 µL from MCS3 and 14 µL from GC2 samples) from these samples in a second GaII Indexing PCR (100 ul reaction, 16 cycles). We combined the initial and supplementary GaII PCR products per sample and concentrated to 15 µL (20 min, 45 °C) using a CentriVap concentrator (Labconco, USA). To clean the PCR products we used AxyPrep beads (1:1.1 PCR products:beads), eluted the beads in 30 µL nuclease-free $H_2O$ and assessed DNA quantity and quality through TapeStation. We prepared an equimolar (6 nM) sequencing pool from all samples, which we concentrated using CentiVap (45 min, 45 °C) to 110 µL, and cleaned using AxyPrep beads (1:1.1 sequencing pool:beads). Following DNA quantity and quality assessment using Qubit, TapeStation, and Fragment Analyzer, we performed one more AxyPrep clean-up (same ratio). We ran final DNA quantity and quality checks via Fragment Analyser and qPCR, and prepared a final sequencing pool (mean fragment size 225 bp, 2.75 nM) for submission to Illumina sequencing (HiSeq XTen, 2 x 150bp cycle) to the KCCG, Darlinghurst Australia.

### ***Bioinformatics***

*Raw sequencing data processing*

Bioinformatic processing and filtering of the sequencing data, hereafter referred to as datasets “Shotgun”, “Phytobaits1” and “HABbaits1”, followed established protocols previously described in detail in Armbrecht et al. (2020). We ran FastQC and MultiQC quality controls on the raw sequencing data and after each step of data filtering (FastQC v0.11.8, Babraham Bioinformatics; MultiQC v1.0.dev0, Ewels et al., 2016). Demultiplexing and adapter trimming was performed using AdapterRemoval v2.3.0 (Schubert et al., 2016), which included the removal of consecutive stretches of low-quality bp and N’s, allowing for a barcode mismatch of 1 bp, discarding reads <25 bp and merging (collapsing) reads into .gz output files. We removed low-complexity sequences using the software Komplexity (--threshold 0.55; Clarke et al., 2019) and removed duplicate sequences using the dedupe tool in BBMap v37.36. To retain the maximum number of reads (especially important for DNA damage assessment), we avoided subsampling (rarefying) our data and used relative abundances in the downstream analyses. We used the NCBI Nucleotide database (ftp://ftp.ncbi.nlm.nih.gov/blast/db/FASTA/nt.gz, downloaded November, 2019) as the reference database to build a MALT index (Step 3) and aligned our sequences using MALT (version 0.4.0; semiglobal alignment) (Herbig et al., 2016). The resulting .blastn files were converted to .rma6 format using the Blast2RMA tool in MEGAN6 (version 6_18_9; Huson et al., 2016) with the default settings except for a minimum support percent of zero (‘off’) and a minimum percent identity of 95%. Subtractive filtering (i.e., subtracting reads for species identified in EBCs from samples) was conducted for each dataset separately following Armbrecht et al. (2020); hereafter, the term ‘samples’ refers to sediment-derived data post-EBC subtraction). Computer code is given in Armbrecht et al., 2020, with the updated program versions used in this publication as per this section.

#### sed*aDNA authenticity assessment and damage analysis*

To assess the authenticity of our Shotgun, Phytobaits1 and HABbaits1 *sed*aDNA we ran the 'MALTExtract' and 'Postprocessing' tools of the HOPS v0.33-2 pipeline (Hübler et al., 2020), including the NCBI mapping and NCBI tree files (13 Nov 2019) provided (https://github.com/rhuebler/HOPS/tree/external/Resources). Configurations deviating from the default HOPS settings included topMaltEx=0.10, minPIdent=95, meganSummary=1, and destackingOff=1.

### References

Armbrecht, L., Herrando-Pérez, S., Eisenhofer, R., Hallegraeff, G.M., Bolch, C.J. and Cooper, A., 2020. An optimized method for the extraction of ancient eukaryote DNA from marine sediments. *Molecular Ecology Resources* 20, 906-919.

Brotherton, P., Haak, W., Templeton, J., Brandt, G., Soubrier, J., Adler, C.J., Richards, S.M., Der Sarkissian, C., Ganslmeier, R., Friederich, S. and Dresely, V., 2013. Neolithic mitochondrial haplogroup H genomes and the genetic origins of Europeans. *Nature communications* 4, 1-11.

Clarke, E.L., Taylor, L.J., Zhao, C., Connell, A., Lee, J.J., Fett, B., Bushman, F.D. and Bittinger, K., 2019. Sunbeam: an extensible pipeline for analyzing metagenomic sequencing experiments. *Microbiome* 7, 46.

De Vargas, C., Audic, S., Henry, N., Decelle, J., Mahé, F., Logares, R., Lara, E., Berney, C., Le Bescot, N., Probert, I. and Carmichael, M., 2015. Eukaryotic plankton diversity in the sunlit ocean. Science 348, 1261605, and companion website http://taraoceans.sb-roscoff.fr/EukDiv/.

Ewels, P., Magnusson, M., Lundin, S., & Käller, M. (2016). multiqc: Summarize analysis results for multiple tools and samples in a single report. Bioinformatics, 32, 3047–3048.

Herbig, A., Maixner, F., Bos, K. I., Zink, A., Krause, J., & Huson, D. H. (2016). malt: Fast alignment and analysis of metagenomic DNA sequence data applied to the Tyrolean Iceman. *BioRxiv*, 050559.

Hübler, R., Key, F.M., Warinner, C., Bos, K.I., Krause, J. and Herbig, A., 2019. HOPS: Automated detection and authentication of pathogen DNA in archaeological remains. *Genome Biology* 20, 1-13.

Huson, D. H., Beier, S., Flade, I., Górska, A., El-Hadidi, M., Mitra, S., & Tappu, R., 2016. MEGAN Community Edition - Interactive exploration and analysis of large-scale microbiome sequencing data. *PLOS Computational Biology 12*, e1004957.

Meyer, M., and Kircher, M., 2010. Illumina sequencing library preparation for highly multiplexed target capture and sequencing. *Cold Spring Harbour Protocols* 6 pdb.prot5448.

MyBaits® Manual v.4.01 — Hybridization Capture for Targeted NGS, 2018. https://arborbiosci.com/wp-content/uploads/2019/08/myBaits-Manual-v4.pdf

Schubert, M., Lindgreen, S., and Orlando, L., 2016. #AdapterRemoval v2: rapid adapter trimming, identification, and read merging. *BMC Research Notes* 9, 88.

### Supplementary Material Tables

**Supplementary Material Table 1.** Eukaryote taxa determined in extraction blank controls (EBCs) of Shotgun, Phytobaits1 and HABbaits1 and removed from downstream analyses.

| Shotgun | SG_21980A_EBC | SG_22301A_EBC | SG_22302A_EBC | SG_23351A_EBC | SG_23352A_EBC | SG_24029A_EBC | SG_24030A_EBC |
|---|---|---|---|---|---|---|---|
| *Plasmodium vivax* | 0 | 0 | 0 | 0 | 2 | 0 | 0 |
| *Plasmodium berghei* | 0 | 0 | 0 | 0 | 0 | 1 | 0 |
| *Eimeria acervulina* | 0 | 1 | 3 | 3 | 0 | 0 | 0 |
| *Stenamoeba* | 0 | 0 | 0 | 1 | 0 | 0 | 0 |
| *Venturia effusa* | 0 | 0 | 0 | 1 | 0 | 0 | 0 |
| *Aureobasidium pullulans* | 0 | 0 | 0 | 1 | 0 | 0 | 0 |
| *Alternaria alternata* | 0 | 0 | 0 | 2 | 0 | 0 | 0 |
| *Alternaria arborescens* | 0 | 0 | 0 | 1 | 0 | 0 | 0 |
| *Penicillium chrysogenum* | 0 | 0 | 0 | 1 | 0 | 0 | 0 |
| *Penicillium expansum* | 0 | 0 | 0 | 1 | 0 | 0 | 0 |
| *Thelebolaceae* | 0 | 0 | 0 | 1 | 0 | 0 | 0 |
| *Beauveria bassiana* | 0 | 0 | 0 | 0 | 1 | 0 | 0 |
| *Cordyceps militaris* | 0 | 1 | 0 | 0 | 3 | 0 | 0 |
| *Lecanicillium* | 0 | 0 | 0 | 0 | 3 | 0 | 0 |
| *Trichoderma* | 0 | 2 | 0 | 0 | 0 | 0 | 0 |
| *Fusarium fujikuroi* | 0 | 0 | 0 | 1 | 0 | 0 | 0 |
| *Fusarium verticillioides* | 0 | 0 | 0 | 7 | 0 | 0 | 0 |
| *Fusarium oxysporum* | 1 | 0 | 0 | 0 | 0 | 0 | 0 |
| *Fusarium graminearum* | 0 | 0 | 0 | 2 | 0 | 0 | 0 |
| *Fusarium pseudograminearum* | 0 | 0 | 0 | 1 | 0 | 0 | 0 |
| *Chaetomium globosum* | 0 | 0 | 0 | 1 | 0 | 0 | 0 |
| *Phaeoacremonium minimum* | 0 | 0 | 0 | 2 | 0 | 0 | 0 |
| *Debaryomyces hansenii* | 2 | 3 | 0 | 0 | 1 | 0 | 0 |
| *Saccharomyces cerevisiae* | 0 | 1 | 0 | 0 | 0 | 0 | 0 |
| *Trametes versicolor* | 0 | 0 | 0 | 2 | 0 | 0 | 0 |
| *Stereum hirsutum* | 0 | 0 | 0 | 12 | 0 | 0 | 0 |
| *Xylobolus sp. 101 OA-2013* | 0 | 0 | 0 | 2 | 0 | 0 | 0 |
| *Filobasidium uniguttulatum* | 0 | 0 | 0 | 1 | 0 | 0 | 0 |
| *Cryptococcus amylolentus* | 0 | 0 | 0 | 1 | 0 | 0 | 0 |
| *Kockovaella* | 0 | 0 | 0 | 1 | 0 | 0 | 0 |
| *Trichosporon asahii* | 1 | 0 | 0 | 1 | 0 | 0 | 0 |
| *Sporidiobolaceae* | 0 | 0 | 0 | 0 | 0 | 0 | 1 |

| | | | | | | | |
|---|---|---|---|---|---|---|---|
| *Sporisorium graminicola* | 0 | 0 | 0 | 1 | 0 | 0 | 0 |
| *Fungi incertae sedis* | 0 | 0 | 0 | 1 | 0 | 0 | 0 |
| *Larimichthys crocea* | 1 | 5 | 0 | 0 | 1 | 0 | 0 |
| *Salarias fasciatus* | 0 | 1 | 0 | 0 | 0 | 2 | 0 |
| *Oncorhynchus nerka* | 0 | 0 | 0 | 0 | 1 | 0 | 0 |
| *Oncorhynchus tshawytscha* | 0 | 0 | 0 | 29 | 0 | 0 | 0 |
| *Clupea harengus* | 0 | 2 | 0 | 0 | 0 | 0 | 0 |
| *Cyprinus carpio* | 25 | 570 | 10 | 41 | 127 | 123 | 27 |
| *Sinocyclocheilus anshuiensis* | 0 | 1 | 0 | 0 | 0 | 0 | 0 |
| *Heterocephalus glaber* | 22 | 5 | 3 | 5 | 1 | 1 | 0 |
| *Ictidomys tridecemlineatus* | 315 | 243 | 38 | 4 | 192 | 51 | 4 |
| *Macaca fascicularis* | 0 | 1 | 0 | 0 | 0 | 0 | 0 |
| *Homo sapiens* | 9 | 17 | 0 | 9 | 21 | 0 | 126 |
| *Pan troglodytes* | 0 | 0 | 0 | 1 | 0 | 0 | 3 |
| *Pongo abelii* | 0 | 15 | 0 | 0 | 0 | 14 | 0 |
| *Aotus nancymaae* | 0 | 0 | 0 | 0 | 0 | 0 | 1 |
| *Panthera pardus* | 0 | 1 | 0 | 0 | 0 | 1 | 0 |
| *Bos mutus* | 0 | 4 | 0 | 0 | 1 | 0 | 0 |
| *Bos taurus* | 3 | 0 | 0 | 0 | 0 | 0 | 0 |
| *Capra hircus* | 0 | 0 | 0 | 1 | 0 | 0 | 0 |
| *Ovis canadensis* | 0 | 1 | 0 | 0 | 0 | 0 | 0 |
| *Odocoileus hemionus* | 0 | 1 | 0 | 0 | 0 | 0 | 0 |
| *Odocoileus virginianus* | 0 | 1 | 0 | 0 | 0 | 0 | 0 |
| *Camelus ferus* | 13 | 7 | 0 | 6 | 0 | 8 | 0 |
| *Aquila chrysaetos* | 0 | 0 | 0 | 33 | 0 | 0 | 0 |
| *Columba livia* | 0 | 0 | 0 | 19 | 0 | 0 | 0 |
| *Streptopelia turtur* | 0 | 0 | 0 | 115 | 0 | 0 | 0 |
| *Falco peregrinus* | 0 | 0 | 0 | 1 | 0 | 0 | 0 |
| *Gallus gallus* | 0 | 0 | 0 | 1 | 0 | 0 | 0 |
| *Corvus brachyrhynchos* | 0 | 0 | 0 | 4 | 0 | 0 | 0 |
| *Cyanistes caeruleus* | 0 | 0 | 0 | 2 | 0 | 0 | 0 |
| *Parus major* | 0 | 0 | 0 | 1 | 0 | 0 | 0 |
| *Zonotrichia albicollis* | 0 | 0 | 0 | 1 | 0 | 0 | 0 |
| *Taeniopygia guttata* | 0 | 0 | 0 | 7 | 0 | 0 | 0 |
| *Serinus canaria* | 0 | 0 | 0 | 2 | 0 | 0 | 0 |
| *Sturnus vulgaris* | 0 | 0 | 0 | 34 | 0 | 0 | 0 |
| *Acrocephalus arundinaceus* | 0 | 0 | 0 | 1 | 0 | 0 | 0 |
| *Phaethon lepturus* | 0 | 0 | 0 | 1 | 0 | 0 | 0 |

| | | | | | | | |
|---|---|---|---|---|---|---|---|
| *Apteryx australis* | 0 | 4 | 0 | 1 | 0 | 0 | 0 |
| *Diphyllobothrium* | 0 | 325 | 0 | 3 | 2 | 35 | 0 |
| *Dicrocoelium dendriticum* | 0 | 8 | 0 | 58 | 0 | 2 | 0 |
| *Trichobilharzia regenti* | 0 | 0 | 0 | 0 | 1 | 0 | 0 |
| *Brugia timori* | 0 | 0 | 0 | 6 | 0 | 0 | 0 |
| *Strongyloides stercoralis* | 0 | 0 | 0 | 0 | 2 | 0 | 0 |
| *Strongyloides venezuelensis* | 0 | 0 | 0 | 1 | 0 | 0 | 0 |
| *Nippostrongylus brasiliensis* | 0 | 2 | 0 | 14 | 39 | 7 | 2 |
| *Parasteatoda tepidariorum* | 0 | 1 | 0 | 0 | 0 | 0 | 0 |
| *Moina brachiata* | 0 | 0 | 0 | 0 | 0 | 1 | 0 |
| *Hyalella azteca* | 0 | 0 | 0 | 2 | 0 | 0 | 0 |
| *Proasellus solanasi* | 0 | 3 | 0 | 1 | 0 | 4 | 0 |
| *Ostrinia furnacalis* | 0 | 0 | 0 | 1 | 0 | 0 | 0 |
| *Harmonia axyridis* | 0 | 0 | 0 | 2 | 0 | 0 | 0 |
| *Drosophila biarmipes* | 0 | 0 | 0 | 1 | 0 | 0 | 0 |
| *Drosophila pseudoobscura* | 0 | 2 | 0 | 0 | 0 | 3 | 0 |
| *Culex pipiens* | 0 | 0 | 0 | 1 | 0 | 0 | 0 |
| *Camponotus floridanus* | 0 | 8 | 0 | 0 | 0 | 0 | 0 |
| *Rhopalosiphum maidis* | 0 | 0 | 0 | 2 | 0 | 0 | 0 |
| *Psylloidea* | 0 | 0 | 0 | 1 | 0 | 0 | 0 |
| *Conus episcopatus* | 0 | 0 | 0 | 1 | 0 | 0 | 0 |
| *Acropora digitifera* | 0 | 25 | 5 | 0 | 0 | 0 | 0 |
| *Dendronephthya gigantea* | 0 | 0 | 0 | 1 | 0 | 0 | 0 |
| *Cercomonadida* | 0 | 0 | 0 | 1 | 0 | 0 | 0 |
| *Saprolegnia parasitica* | 0 | 0 | 0 | 1 | 0 | 0 | 0 |
| *Chlorococcum tatrense* | 0 | 0 | 0 | 4 | 0 | 0 | 0 |
| *Volvox carteri* | 0 | 0 | 0 | 1 | 0 | 0 | 0 |
| *Mychonastes homosphaera* | 0 | 0 | 0 | 2 | 0 | 0 | 0 |
| *Monoraphidium neglectum* | 0 | 0 | 0 | 1 | 0 | 0 | 0 |
| *Micractinium conductrix* | 0 | 0 | 0 | 1 | 0 | 0 | 0 |
| *Coccomyxa sp. SUA001* | 0 | 0 | 0 | 1 | 0 | 0 | 0 |
| *Physcomitrella patens* | 1 | 0 | 0 | 0 | 0 | 0 | 0 |
| *Dioon* | 9 | 38 | 0 | 3 | 0 | 89 | 0 |
| *Pinus taeda* | 0 | 0 | 0 | 8 | 0 | 0 | 0 |
| *Daucus carota* | 0 | 0 | 0 | 12 | 0 | 0 | 0 |
| *Lasthenia californica* | 22 | 0 | 0 | 0 | 0 | 0 | 0 |
| *Chionanthus rupicola* | 0 | 0 | 0 | 1 | 0 | 0 | 0 |

| | | | | | | | |
|---|---|---|---|---|---|---|---|
| *Physochlaina orientalis* | 0 | 0 | 0 | 1 | 0 | 0 | 0 |
| *Nepenthes ventricosa x N. alata* | 0 | 0 | 0 | 1 | 0 | 0 | 0 |
| *Phaseolus* | 0 | 1 | 0 | 0 | 0 | 0 | 0 |
| *Quercus suber* | 0 | 0 | 0 | 2 | 0 | 0 | 0 |
| *Populus trichocarpa* | 0 | 0 | 0 | 3 | 0 | 0 | 0 |
| *Pyrus x bretschneideri* | 0 | 0 | 0 | 0 | 0 | 0 | 1 |
| *Theobroma cacao* | 0 | 5 | 0 | 0 | 0 | 0 | 0 |
| *Rhodamnia argentea* | 0 | 1 | 0 | 0 | 0 | 0 | 0 |
| *Citrus sinensis* | 0 | 0 | 0 | 1 | 0 | 0 | 0 |
| *Elaeis guineensis* | 0 | 0 | 0 | 5 | 0 | 0 | 0 |
| *Oryza sativa* | 0 | 0 | 0 | 1 | 0 | 0 | 0 |
| *Triticum aestivum* | 0 | 1 | 0 | 0 | 0 | 0 | 0 |
| *Triticum monococcum* | 0 | 0 | 0 | 0 | 0 | 0 | 1 |
| *Zea mays* | 0 | 0 | 0 | 3 | 0 | 0 | 0 |

| **Phytobaits1** | **HYB18SV9_A21980_EBC** | **HYB18SV9_A22301_EBC** | **HYB18SV9_A22302_EBC** | **HYB18SV9_A23351_EBC** | **HYB18SV9_A23352_EBC** | **HYB18SV9_A24029_EBC** | **HYB18SV9_A24030_EBC** |
|---|---|---|---|---|---|---|---|
| *Opegrapha vulgata* | 0 | 0 | 0 | 0 | 0 | 0 | 1 |
| *Aureobasidium pullulans* | 0 | 0 | 0 | 2 | 0 | 0 | 0 |
| *Alternaria solani* | 0 | 0 | 0 | 1 | 0 | 0 | 0 |
| *Penicillium* | 0 | 0 | 0 | 2 | 0 | 0 | 0 |
| *Metarhizium robertsii* | 0 | 0 | 0 | 0 | 0 | 0 | 1 |
| *Fusarium graminearum* | 0 | 0 | 0 | 1 | 0 | 0 | 0 |
| *Microascales* | 0 | 0 | 0 | 1 | 0 | 0 | 0 |
| *Saccharomyces cerevisiae* | 0 | 1 | 0 | 0 | 0 | 0 | 0 |
| *Stereum hirsutum* | 0 | 0 | 0 | 5 | 0 | 0 | 0 |
| *Cryptococcus neoformans* | 0 | 0 | 0 | 0 | 0 | 0 | 44 |
| *Kockovaella* | 0 | 0 | 0 | 3 | 0 | 0 | 0 |
| *Sporidiobolaceae* | 0 | 0 | 0 | 0 | 0 | 0 | 1 |
| *Malassezia restricta* | 0 | 0 | 0 | 0 | 0 | 0 | 2 |
| *Sporisorium graminicola* | 0 | 0 | 0 | 1 | 0 | 0 | 0 |
| *Mucorales* | 0 | 0 | 0 | 1 | 0 | 0 | 0 |
| *Salarias fasciatus* | 0 | 0 | 1 | 0 | 0 | 1 | 0 |
| *Oncorhynchus tshawytscha* | 0 | 0 | 0 | 7 | 0 | 0 | 0 |
| *Cyprinus carpio* | 0 | 0 | 4 | 0 | 8 | 51 | 4 |
| *Heterocephalus glaber* | 0 | 0 | 1 | 0 | 0 | 0 | 0 |
| *Ictidomys tridecemlineatus* | 1 | 4 | 19 | 0 | 1 | 28 | 1 |
| *Macaca fascicularis* | 0 | 0 | 0 | 0 | 0 | 0 | 1 |

| | | | | | | | |
|---|---|---|---|---|---|---|---|
| *Theropithecus gelada* | 0 | 0 | 0 | 0 | 0 | 0 | 1 |
| *Homo sapiens* | 1 | 3 | 0 | 5 | 3 | 1 | 650 |
| *Pan troglodytes* | 0 | 0 | 0 | 0 | 0 | 0 | 17 |
| *Pongo abelii* | 0 | 0 | 0 | 0 | 0 | 5 | 1 |
| *Vulpes vulpes* | 0 | 0 | 0 | 0 | 0 | 0 | 1 |
| *Lynx canadensis* | 0 | 0 | 0 | 0 | 0 | 0 | 1 |
| *Ovis canadensis* | 0 | 1 | 0 | 0 | 0 | 0 | 0 |
| *Odocoileus* | 0 | 2 | 0 | 0 | 0 | 0 | 0 |
| *Camelus ferus* | 0 | 0 | 0 | 0 | 0 | 2 | 0 |
| *Phyllostomidae* | 0 | 0 | 0 | 1 | 0 | 0 | 0 |
| *Aquila chrysaetos* | 0 | 0 | 0 | 8 | 0 | 0 | 0 |
| *Columba livia* | 0 | 0 | 0 | 3 | 0 | 0 | 0 |
| *Streptopelia turtur* | 0 | 0 | 0 | 29 | 0 | 0 | 0 |
| *Gallus gallus* | 0 | 0 | 1 | 0 | 0 | 0 | 0 |
| *Cyanistes caeruleus* | 0 | 0 | 0 | 1 | 0 | 0 | 0 |
| *Sturnus vulgaris* | 0 | 0 | 0 | 5 | 0 | 0 | 0 |
| *Geospiza fortis* | 0 | 0 | 0 | 1 | 0 | 0 | 0 |
| *Diphyllobothrium* | 0 | 3 | 0 | 0 | 0 | 84 | 0 |
| *Spirometra erinaceieuropaei* | 0 | 0 | 0 | 0 | 0 | 0 | 1 |
| *Brugia timori* | 0 | 0 | 0 | 1 | 0 | 0 | 0 |
| *Onchocerca ochengi* | 0 | 0 | 0 | 1 | 0 | 0 | 0 |
| *Nippostrongylus brasiliensis* | 0 | 0 | 0 | 0 | 1 | 0 | 0 |
| *Proasellus solanasi* | 0 | 0 | 0 | 0 | 0 | 1 | 0 |
| *Scaptodrosophila lebanonensis* | 0 | 0 | 0 | 0 | 0 | 1 | 0 |
| *Drosophila pseudoobscura* | 0 | 0 | 0 | 0 | 0 | 1 | 0 |
| *Sciaroidea* | 0 | 0 | 0 | 1 | 0 | 0 | 0 |
| *Culex pipiens* | 0 | 0 | 0 | 2 | 0 | 0 | 0 |
| *Camponotus floridanus* | 0 | 2 | 0 | 0 | 0 | 0 | 0 |
| *Diaspididae* | 0 | 0 | 0 | 1 | 0 | 0 | 0 |
| *Pontoscolex corethrurus* | 0 | 0 | 0 | 0 | 0 | 1 | 0 |
| *Cercomonadida* | 0 | 0 | 0 | 1 | 0 | 0 | 0 |
| *Stramenopiles* | 0 | 0 | 0 | 6 | 0 | 0 | 0 |
| *Volvox carteri* | 0 | 0 | 0 | 1 | 0 | 0 | 0 |
| *Micractinium conductrix* | 0 | 0 | 0 | 1 | 0 | 0 | 0 |
| *Coccomyxa subellipsoidea* | 0 | 0 | 0 | 1 | 0 | 0 | 0 |
| *Physcomitrella patens* | 0 | 0 | 0 | 0 | 0 | 1 | 0 |
| *Dioon* | 0 | 0 | 0 | 0 | 0 | 94 | 0 |
| *Pinus <subgenus>* | 0 | 0 | 0 | 1 | 0 | 0 | 0 |
| *Daucus carota* | 0 | 0 | 0 | 3 | 0 | 0 | 0 |

| | | | | | | | |
|---|---|---|---|---|---|---|---|
| *Lasthenia californica* | 0 | 0 | 0 | 0 | 0 | 3 | 0 |
| *Oreocharis mileensis* | 0 | 0 | 0 | 1 | 0 | 0 | 0 |
| *Fagales* | 0 | 0 | 0 | 14 | 0 | 0 | 0 |
| *Populus trichocarpa* | 0 | 0 | 0 | 2 | 0 | 0 | 0 |
| *Malus domestica* | 0 | 0 | 0 | 0 | 0 | 0 | 2 |
| *Oryza sativa* | 0 | 0 | 0 | 1 | 0 | 0 | 0 |
| *Triticum aestivum* | 0 | 0 | 0 | 0 | 0 | 0 | 1 |
| *Zea mays* | 0 | 0 | 0 | 2 | 0 | 0 | 0 |
| *Dioscorea rotundata* | 0 | 0 | 0 | 1 | 0 | 0 | 0 |

| **HABbaits1** | **HYBHAB_A21980_EBC** | **HYBHAB_A22301_EBC** | **HYBHAB_A22302_EBC** | **HYBHAB_A23351_EBC** | **HYBHAB_A23352_EBC** | **HYBHAB_A24029_EBC** | **HYBHAB_A24030_EBC** |
|---|---|---|---|---|---|---|---|
| *Stenamoeba* | 0 | 0 | 0 | 1 | 0 | 0 | 0 |
| *Opegrapha vulgata* | 0 | 0 | 0 | 0 | 0 | 0 | 1 |
| *Pleosporales* | 0 | 0 | 0 | 1 | 0 | 0 | 0 |
| *Eurotiales* | 0 | 0 | 0 | 1 | 0 | 0 | 0 |
| *Agaricomycetes incertae sedis* | 0 | 0 | 0 | 1 | 0 | 0 | 0 |
| *Malassezia restricta* | 0 | 0 | 0 | 0 | 0 | 0 | 1 |
| *Rhizophydiales* | 0 | 0 | 0 | 1 | 0 | 0 | 0 |
| *Cyprinus carpio* | 0 | 0 | 0 | 0 | 3 | 2 | 1 |
| *Ictidomys tridecemlineatus* | 0 | 0 | 3 | 0 | 0 | 0 | 0 |
| *Theropithecus gelada* | 0 | 0 | 0 | 0 | 0 | 0 | 1 |
| *Homo sapiens* | 0 | 0 | 0 | 1 | 0 | 0 | 83 |
| *Pongo abelii* | 0 | 0 | 0 | 0 | 0 | 1 | 0 |
| *Lynx canadensis* | 0 | 0 | 0 | 0 | 0 | 0 | 1 |
| *Aquila chrysaetos* | 0 | 0 | 0 | 1 | 0 | 0 | 0 |
| *Streptopelia turtur* | 0 | 0 | 0 | 1 | 0 | 0 | 0 |
| *Corvus brachyrhynchos* | 0 | 0 | 0 | 1 | 0 | 0 | 0 |
| *Zonotrichia albicollis* | 0 | 0 | 0 | 1 | 0 | 0 | 0 |
| *Diphyllobothrium* | 0 | 1 | 0 | 0 | 0 | 3 | 0 |
| *Rhabditidae* | 0 | 0 | 0 | 1 | 0 | 0 | 0 |
| *Harmonia axyridis* | 0 | 0 | 0 | 1 | 0 | 0 | 0 |
| *Sciaroidea* | 0 | 0 | 0 | 1 | 0 | 0 | 0 |
| *Diaspididae* | 0 | 0 | 0 | 1 | 0 | 0 | 0 |
| *Cercozoa* | 0 | 0 | 0 | 2 | 0 | 0 | 0 |
| *Stramenopiles* | 0 | 0 | 0 | 1 | 0 | 0 | 0 |
| *Chlorophyta* | 0 | 0 | 0 | 1 | 0 | 0 | 0 |
| *Dioon* | 0 | 0 | 0 | 0 | 0 | 6 | 0 |
| *Malus domestica* | 0 | 0 | 0 | 0 | 0 | 0 | 2 |
| *Triticum aestivum* | 0 | 0 | 0 | 0 | 0 | 0 | 1 |
